# Human-specific lncRNA TMEM9B-AS1 is downregulated in skeletal muscle of individuals with type 2 diabetes and regulates ribosomal biogenesis

**DOI:** 10.1101/2024.07.05.602204

**Authors:** Ilke Sen, Jonathon A B Smith, Elena Caria, Mladen Savikj, Kristian Lian, Stian Ellefsen, Juleen R Zierath, Anna Krook

## Abstract

Long non-coding RNAs (lncRNAs) are important regulators of skeletal muscle physiology, with altered expression noted in several human diseases including type 2 diabetes. Here we report TMEM9B-AS1, a previously uncharacterized lncRNA, is downregulated in skeletal muscle of men with type 2 diabetes. Silencing of TMEM9B-AS1 in primary human myotubes attenuated protein synthesis, concomitant with reduced phosphorylation of ribosomal protein S6, a downstream target of ERK and mTOR pathways. Moreover, we provide evidence that TMEM9B-AS1 plays a pivotal role in the regulation of ribosomal biogenesis by facilitating mRNA stabilization of the transcription factor MYC through a direct physical interaction with the RNA-binding protein IGF2BP1. Disrupted ribosomal biogenesis resulting from TMEM9B-AS1 silencing is linked reduced skeletal muscle mass in type 2 diabetes, elucidating molecular mechanisms contributing to skeletal muscle loss in metabolic disease.

## INTRODUCTION

Long non-coding RNAs (lncRNAs) make up a considerable part of the genome and encompass a diverse class of RNA molecules that are not translated into proteins ^1^. As important regulators of cell and tissue function^2^ lncRNAs have been implicated in the development of metabolic diseases. Type 2 diabetes is a metabolic disease exacting an ever-increasing toll on public health. The aetiology of type 2 diabetes is complex, involving multiple tissues resulting in impaired glucose homeostasis. While several lncRNAs have been shown to regulate glucose and lipid metabolism in liver and adipose tissue ^3,4^, as well as functional properties of pancreatic islets ^3^, effects in skeletal muscle are not clearly understood. Skeletal muscle comprises ∼50% of body mass, and as the primary site for insulin-stimulated glucose disposal, impaired insulin action in this organ often presents early in type 2 diabetes pathogenesis ^5^. Furthermore, type 2 diabetes is associated with a decline in muscle mass, strength, and function ^6^, which can be exacerbated by GLP-1 receptor agonists prescribed as weight loss therapy.

Functional roles attributed to lncRNAs include the regulation of transcriptional and post-translational processes ^4^. Cytoplasmic lncRNAs interact with RNA-binding proteins to impact mRNA stability of specific genes and thereby coordinate protein synthesis/translation ^7^. Protein synthesis rates result from the efficiency and capacity of the translation machinery. As total ribosomal content dictates translational capacity, ribosomal biogenesis is important for the maintenance and hypertrophy of skeletal muscle mass ^8^. Ribosome biogenesis is modulated by several signalling pathways, with ncRNAs emerging as notable regulators of this process^9^. Although the precise mechanisms remain unclear, the PI3K/mTOR/p70S6K1 network and the transcription factor MYC appear necessary for ribosomal biogenesis in skeletal muscle ^10^.

In a search for lncRNA’s associated with insulin resistance, we identified lncRNA TMEM9B-AS1 expression is downregulated in skeletal muscle of men with type 2 diabetes. Contextualizing the role of this differentially expressed lncRNA in metabolic processes, we provide evidence that TMEM9B-AS1 is required for protein synthesis and regulates ribosomal biogenesis by facilitating the stability of MYC mRNA through physical interaction with the RNA-binding protein IGF2BP1. This interaction is essential for ribosomal biogenesis and protein synthesis in myocytes. Thus, TMEM9B-AS1 is a potential target to improve metabolic health and physical performance by ameliorating detriments in skeletal muscle mass and function.

## Results

### TMEM9B-AS1 is downregulated in skeletal muscle of men with type 2 diabetes and is localized to the cytoplasm in primary human myotubes

Differentially expressed transcripts were determined from skeletal muscle biopsies obtained from a previously published cohort of 19 men with type 2 diabetes and 17 men with normal glucose tolerance ^11^ (Fig. 1a). TMEM9B-AS1, a human specific antisense lncRNA, was downregulated in skeletal muscle of individuals with type 2 diabetes. Reduced TMEM9B-AS1 expression was validated in a sub-group of individuals with type 2 diabetes and normal glucose tolerance from a separate cohort^12^ (Fig. 1b). Bulk expression of TMEM9B-AS1 in different human tissues from both females and males was probed in the GTEX database (Fig. 1c) revealing broad tissue distribution of TMEM9B-AS1 expression (Fig. 1d). Since several of the participants with type 2 diabetes were treated with Metformin or statins, we determined whether these drugs would directly alter TMEM9B-AS1 expression. TMEM9B-AS1 levels were unaltered following a 3-hour exposure to Metformin (5 µM, 200 µM, or 2 mM) or Simvastatin (5 µM) (Extended Data Fig 1). Thus, reduced TMEM9B-AS1 levels appear to be a feature of type 2 diabetes rather than a response to Metformin or Simvastatin treatment.

**Figure 1.**
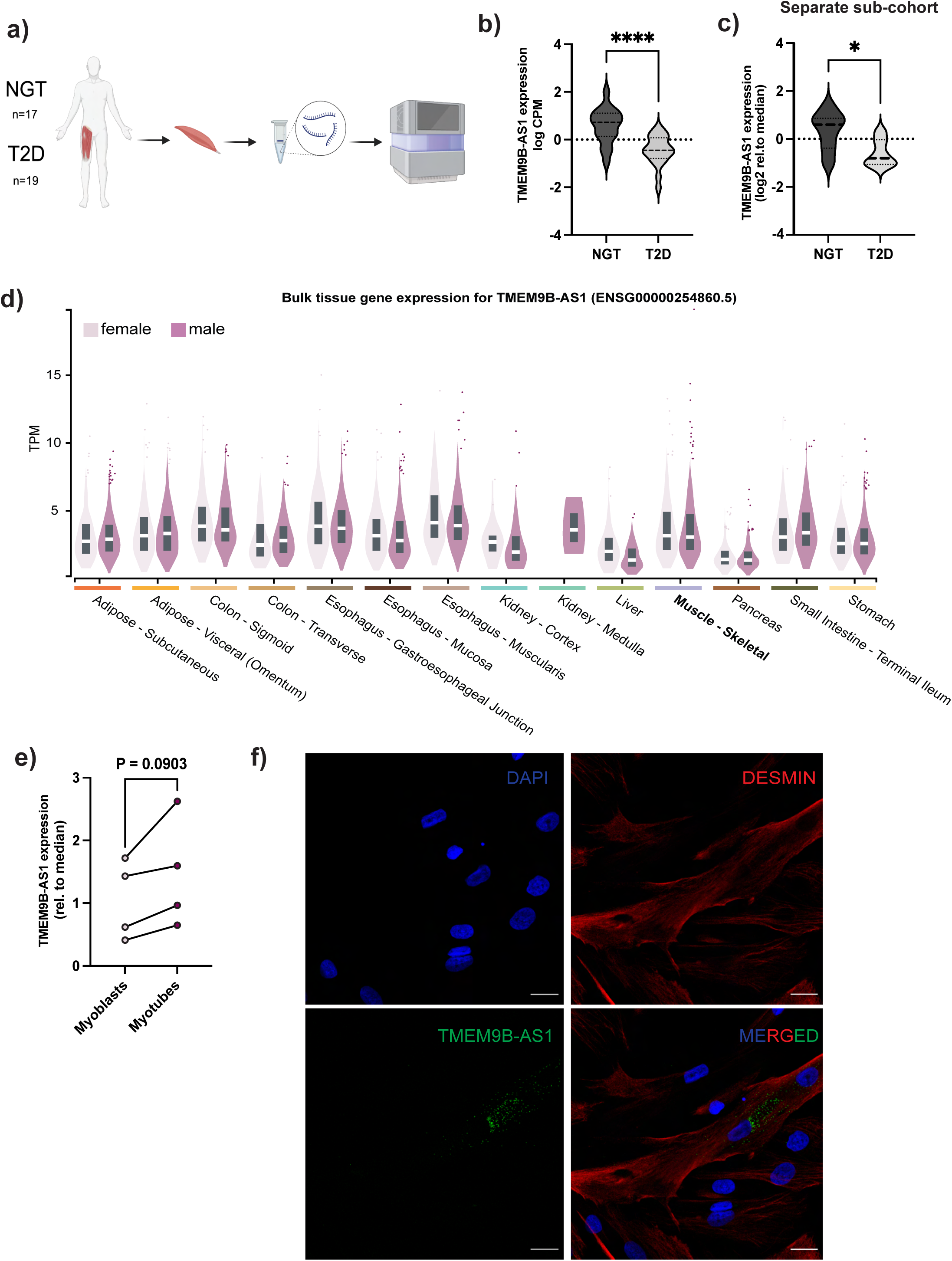
TMEM9B-AS1 is downregulated in skeletal muscle from people with type 2 diabetes and expressed in the cytoplasm of cultured human myotubes. **a.** Schematic representation of the study design. **b.** RNA-SEQ data showing a decrease in TMEM9B-AS1 expression levels in skeletal muscle of the people with type 2 diabetes (n=19) compared to people with normal glucose tolerance (n=17). **** p<0.0001. **c.** Gene expression analysis with Taqman assays in a separate sub-cohort of people with type 2 diabetes (n=6) and matched normal glucose tolerant people (n=7) * p<0.05. **d.** Bulk tissue gene expression levels of TMEM9B-AS1 in several metabolic tissues from both females and males. Data was extracted from the GTEX database. **e.** Gene expression analysis with Taqman assays showing the expression of TMEM9B-AS1 in human skeletal muscle myoblasts and differentiated myotubes (n=4). **f.** Confocal images showing the localization of TMEM9B-AS1 by RNA FISH combined with immunofluroscence. Desmin antibody was used as a structural protein marker for human myotubes and DAPI was used to stain the nucleus. Scale bar is 20 µm.

Skeletal muscle is a complex heterogeneous tissue that consists of multinucleated muscle fibres, endothelial cells, satellite cells (muscle stem cells), immune cells and other mononuclear cells ^13^. To investigate whether TMEM9B-AS1 is expressed in myocytes, we determined TMEM9B-AS1 expression using QPCR in primary human skeletal muscle cells. TMEM9B-AS1 was expressed in human myoblasts as well as differentiated human myotubes (Fig. 1e). To determine the subcellular localization of TMEM9B-AS1 we used RNA-FISH combined with immunocytochemistry (Fig. 1f). Human myotubes were labelled using a specific antibody against Desmin (DES), a muscle-specific protein important for myotube structure, and a specific probe to detect TMEM9B-AS1 RNA. TMEM9B-AS1 is localised in the cytoplasm, close to the nuclear membrane where the ribosome enriched endoplasmic reticulum is located (Fig. 1f).

### TMEM9B-AS1 regulates protein synthesis through mTOR and ERK signalling

To investigate the role of TMEM9B-AS1 in skeletal muscle metabolism we silenced TMEM9B-AS1 expression with siRNA in primary human skeletal muscle cells. TMEM9B-AS1 expression was efficiently reduced following siRNA silencing (Fig. 2a). Abundance of proteins important for muscle differentiation and structure, including Desmin, (DES, Extended Data Fig. 2a), MYH7 (MyHC-I, Extended Data Fig. 2b), MYH1/2 (MyHC-IIX/IIA, Extended Data Fig. 2c) were reduced following TMEM9B-AS1 silencing, with no difference seen for myogenin (MYOG; Extended Data Fig 2d). Furthermore, TMEM9B-AS1 silencing did not affect glucose oxidation (Extended Data Fig. 3a), lipid (palmitic acid) oxidation (Extended Data Fig. 3b) or glucose uptake (Extended Data Fig. 3c) in human myotubes at baseline or following appropriate stimulation. However, decreased protein content was a consistent finding following TMEM9B-AS1 silencing in human myotubes (Fig. 2b). Thus, we postulated that TMEM9B-AS1 may play a role in skeletal muscle protein metabolism, including protein synthesis/translation or protein degradation, which are dynamically regulated processes that orchestrate the gain or loss of muscle mass ^14,15^. To determine whether TMEM9B-AS1 is involved in protein degradation, we assessed effects on the ubiquitin-proteasome system, which is activated during muscle atrophy and may contribute to loss of muscle mass ^15^. Proteins undergo proteasomal degradation in the ubiquitin-proteasome system ^15,16^ following ubiquitination by E3 ubiquitin ligases ^15,17^. Upon siRNA silencing of TMEM9B-AS1, no differences were observed on the total ubiquitination of proteins (Fig. 2c) or the abundance of muscle-specific E3 ligase, atrogin-1/MAFbx (Fig. 2d). In contrast, we detected the reduction of several subunits (1,2,3,5,6,7) of proteasome 20S alpha (Fig. 2e) that are typically elevated in catabolic conditions modulating muscle loss ^17^. Taken together, these findings indicate that the reduction in total protein content was not attributable to protein degradation, thus suggesting the potential regulation of protein synthesis by TMEM9B-AS1.

**Figure 2.**
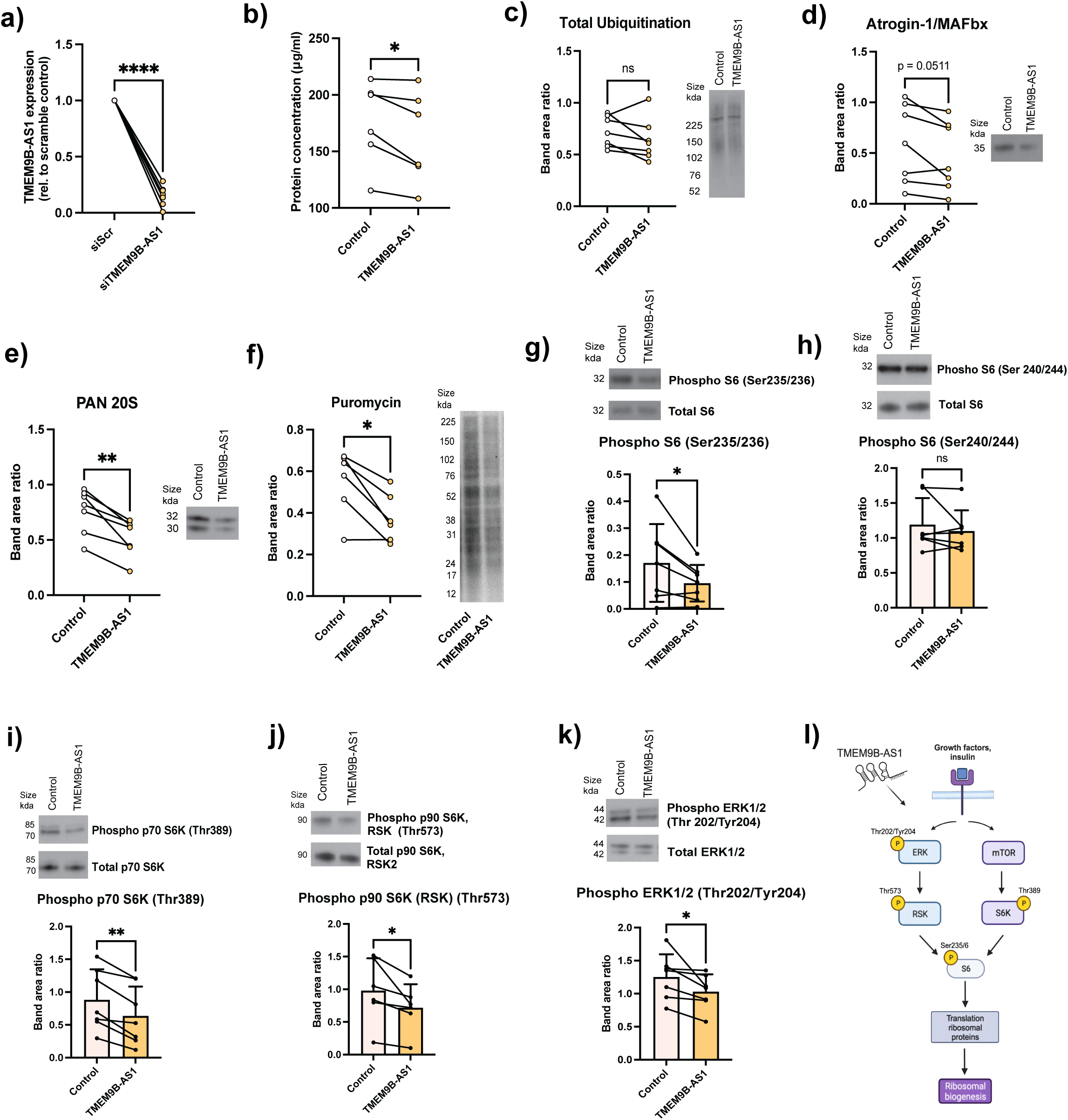
TMEM9B-AS1 regulates protein synthesis through mTOR and ERK signalling in human myotubes. **a.** QPCR results showing silencing efficiency of TMEM9B-AS1 in human myotubes in 7 different donors. **b.** BCA results showing a decrease in total protein levels upon TMEM9B-AS1 silencing, n=6. Western blot results of **c.** Total ubiquitination (n=7), **d.** Atrogin/MAFbx (n=7), **e.** PAN20S (n=7). **f.** Western blot results of puromycin following a Surface sensing of translation (SUnSET) assay, n=6. Western blot results showing the level of phosphorylation of **g.** S6 protein on Ser 235/236 (n=7), **h.** S6 protein on Ser 240/244 (n=7), **i.** p70 S6K on Thr389 (n=7), **j.** p90 S6K (RSK) on Thr573 (n=6), and **k.** ERK1/2 on Thr202/Tyr204 (n=7) upon TMEM9B-AS1 silencing. Phosphorylation levels were normalised to total protein levels following normalisation to a selected ponceau band. For statistical analysis a two-tailed paired t-test was performed, *p<0.05, **p<0.01. **l.** Schematic representation of the regulation of ERK and mTOR pathway and the downstream process by TMEM9B-AS1.

To determine whether TMEM9B-AS1 plays a role in protein synthesis we performed Surface sensing of translation (SUnSET) assay, tracing puromycin incorporation into newly synthesized proteins. Subsequently, we found that TMEM9B-AS1 silencing in human myotubes resulted in a decrease in protein synthesis/translation (Fig. 2f). To understand how TMEM9B-AS1 regulates protein synthesis/translation, we first focused on the mTORC1 pathway as a master regulator of protein synthesis/translation ^15,18^. No changes were noted on mTOR phosphorylation (Extended Data Fig. 4a), or on the downstream target 4E-BP1 (Extended Data Fig. 4b) following TMEM9B-AS1 silencing. However, assessing ribosomal protein S6, representing a separate signalling branch downstream of mTORC1 ^19^, we observed that phosphorylation of ribosomal protein S6 on Ser235/236 (Fig. 3g), but not Ser240/244 (Fig. 3h) was decreased by silencing of TMEM9B-AS1. While phosphorylation of S6 protein on the Ser235/236 site is regulated by two S6Ks: p70 S6K (downstream of mTOR) and p90 S6Ks (RSKs; downstream of ERK signalling), the Ser240/244 phosphorylation site is targeted specifically by p70 S6K^20^. This data suggested that two of the upstream S6-kinases, p70 and p90, could be involved in TMEM9B-AS1-sensitive phospho-regulation of S6 protein. Next, we determined whether loss of TMEM9B-AS1 affected the phosphorylation of p70 and p90 S6-kinases and noted that phosphorylation of p70S6K Thr389 (Fig. 2i) and p90S6K Thr573 (Fig. 2j) were reduced following silencing of TMEM9B-AS1. p90 S6-Kinase is regulated by ERK 1/2, and silencing of TMEM9B-AS1 in human myotubes also decreased ERK1/2 phosphorylation on Thr202/Tyr204 (Fig. 2k). Taken together our data suggests that TMEM9B-AS1 regulates protein synthesis/translation through ERK and mTOR pathways and the phosphorylation of the downstream S6 protein, specifically by modulating the Ser235/236 phosphorylation site that is targeted by p70 and p90 S6Ks (Fig. 2l). Phosphorylation of S6 protein is necessary for translation of ribosomal proteins required for ribosomal biogenesis ^21^, which indicated that TMEM9B-AS1 could be involved in the regulation of ribosomal biogenesis.

**Figure 3.**
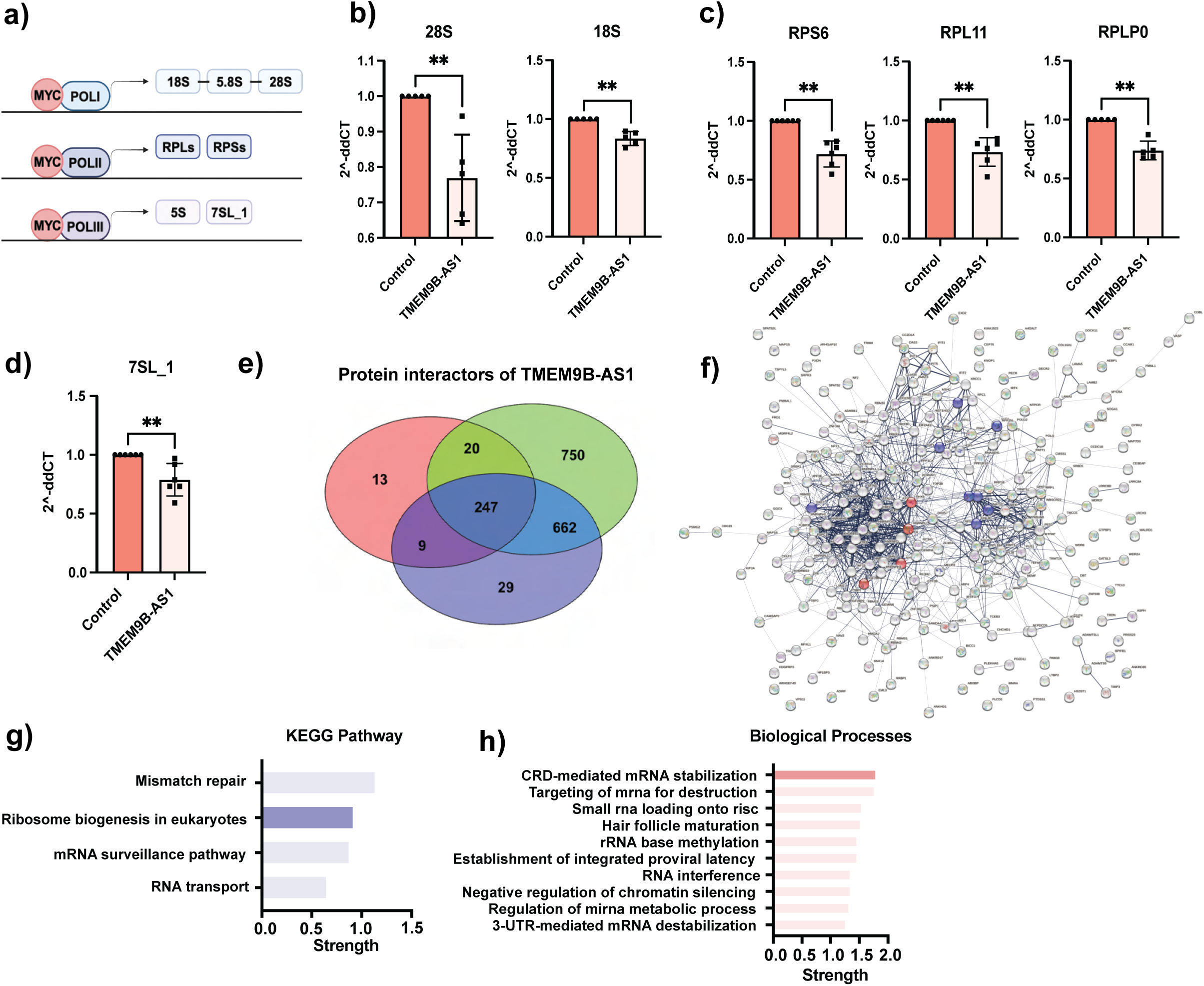
TMEM9B-AS1 regulates ribosomal biogenesis in human myotubes. **a.** Schematic representation of the transcriptional regulation of ribosomal genes/RNAs mediated by different polymerases. **b.** QPCR results showing a decrease in POLI **c.** POLII and **d.** POLIII dependent ribosomal RNA levels upon TMEM9B-AS1 silecing. The geometric means of TBP and GUSB genes were used as reference. n=6, * p<0.05, ** p<0.01. **e.** Results of the RNA pulldown experiment combined with Mass spectroscopy. Venn diagrams show the number of specific binding partners of TMEM9B-AS1 in three different RNA pulldowns and their overlaps. **f.** Protein-protein interaction networks achieved from STRING database. **g.** KEGG pathway and **h.** GO-term Biological Processes functional enrichment analysis conducted on STRING database using specific protein interactors of TMEM9B-AS1 found in all pulldowns. Protein interactors of TMEM9B-AS1 are enriched in pathways related to ribosomal biogenesis. For KEGG pathway, ribosomal biogenesis in eukaryotes, Strength= 0,92, FDR=0,0018. For Biological processes, CRD-mediated mRNA stabilization enrichment term, Strength= 1,8, FDR= 0,00033.

### TMEM9B-AS1 regulates ribosomal biogenesis

*De novo* synthesis of ribosomes requires coordinated synthesis of equimolar quantities of all four types of rRNA subunits and over 80 ribosomal proteins. This, in turn, involves all three polymerases, POLI, POLII and POLIII, each of which interacts with the master transcription factor MYC, serving as a direct regulator of ribosome biogenesis ^19^ (Illustrated in Fig. 3a). To determine the involvement of TMEM9B-AS1 in ribosomal biogenesis, we measured expression of several ribosomal genes/RNAs that depend on POL I (Fig. 3b), POL II (Fig. 3c), and POL III (Fig. 3d) following siRNA silencing of TMEM9B-AS1 in human myotubes. The expression of ribosomal genes/RNAs mediated by all three polymerases were markedly decreased upon reduction of TMEM9B-AS1.

To probe whether TMEM9B-AS1 directly interacts with MYC, we performed an RNA pull-down assay using TMEM9B-AS1 as bait, followed by Western blot analysis for MYC. We report evidence against a direct interaction between TMEM9B-AS1 and MYC. To understand how TMEM9B-AS1 regulates ribosomal biogenesis, we took an unbiased approach to assess potential protein interactions with TMEM9B-AS1 using an RNA pulldown experiment followed by mass spectroscopy (MS). A list of protein interactors of TMEM9B-AS1 was generated by considering the overlapping proteins as specific interactors in three individual pulldowns (Fig. 3e). Using this list, we probed the STRING database for potential protein-protein interaction networks (Fig. 3f) and performed functional enrichment analysis using KEGG pathway enrichment (Fig. 3g) and Go-term Biological Processes (Fig. 3h). We observed that protein interactors of TMEM9B-AS1 were enriched in pathways related to ribosomal biogenesis (Fig 3g). Functional enrichment analysis for Biological Processes revealed that protein interactors of TMEM9B-AS1 were also enriched in Coding region instability determinant (CRD)-mediated mRNA stabilization processes derived by RNA-binding proteins. CRD is a 249-nucleotide region on MYC mRNA that is required for its stabilization ^22^. This data implied that TMEM9B-AS1 could be involved in the regulation of MYC mRNA stabilization. Indeed, in cultured human muscle cells, the subcellular localisation of TMEM9B-AS1 is consistent with a ribosomal localisation (Fig 1f). Furthermore, reanalysis of ribosome profiling data from a previous study ^23^ identifies physical association between TMEM9B-AS1 and ribosomes in several tissues and cell types.

### TMEM9B-AS1 regulates MYC mRNA stabilization by physically interacting with the RNA-binding proteins IGF2BPs

To further determine specific interactors of TMEM9B-AS1, we compared the list of protein interactors in the RNA pulldown-MS data with TMEM9B-AS1 binding proteins identified in publicly available CLIP-SEQ data in the ENCORI/starBase database ^24^. From this confirmation, we generated a short-list of unique interactors of TMEM9B-AS1 (Fig. 4a). Using this short-list of proteins, we performed functional enrichment analysis for the biological processes and again found that the top GO term was CRD-mediated mRNA stabilization, mostly derived from the RNA-binding proteins insulin-like growth factor 2 mRNA binding proteins (IGF2BPs); IGF2BP1 and IGF2BP2 (Fig. 4a-b). IGF2BPs are highly conserved RNA-binding proteins that play a role in regulating RNA processing at different stages, including localization, translation, and stability ^25,26^.

**Figure 4.**
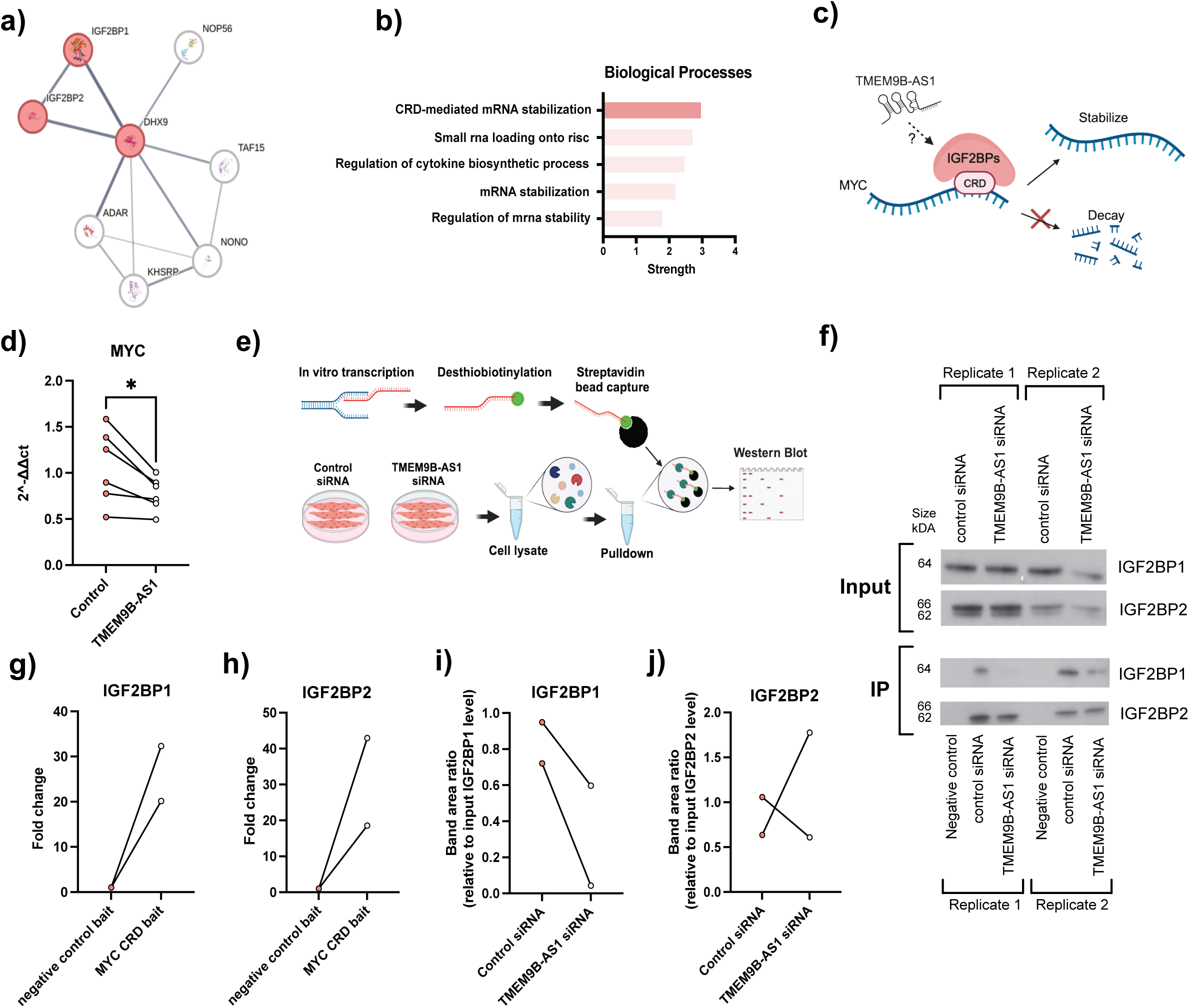
TMEM9B-AS1 regulates MYC mRNA stability through physical interaction with RNA-binding IGF2BP1in human myotubes. **a.** Unique protein interactors of TMEM9B-AS1, determined by using the overlapping proteins in three individual TMEM9B-AS1 RNA pull-downs and TMEM9B-AS1 binding proteins according to the CLIP-SEQ data found in publicly available ENCORI database **b.** Functional enrichment analysis for the biological processes on STRING database using unique protein interactors of TMEM9B-AS1 achieved by the comparison of specific interactors of TMEM9B-AS1 from RNA pulldown data and CLIP-SEQ data from ENCORI database. CRD-mediated mRNA stabilization is the top enriched term. Strength= 2,99, FDR=0,0023**. c.** Schematic representation of interaction between MYC mRNA and RNA-binding proteins, IGF2BPs via the CRD region on MYC mRNA, which facilitates the stabilization of MYC mRNA. **d.** mRNA levels of MYC upon TMEM9B-AS1 silencing. n=6, * p<0.05. **e.** Schematic representation of RNA pull-down experiments to determine the binding efficiency of RNA-binding proteins IGF2BP1 and IGF2BP2 to CRD region on MYC mRNA in the presence or absence of TMEM9B-AS1. **f.** Western blot for IGF2BP1 and IGF2BP2 upon RNA-pulldowns. Negative control RNA was used to determine the RNA pull-down specificity over MYC CRD. RNA pull-down specificity of **g.** IGF2BP1 and **h.** IGF2BP2. RNA binding efficiency of **i.** IGF2BP1 and **j.** IGF2BP2 upon two RNA pull-downs in control and TMEM9B-AS1 silencing conditions. Data were normalized to the individual input levels of IGF2BP1 and IGF2BP2.

IGF2BP1 stabilises MYC mRNA by associating with its CRD region in cell culture models^19,22,26^ (illustrated in Fig 4c), and we hypothesized that TMEM9B-AS1 could play a role in regulating this process. Accordingly, MYC mRNA levels were reduced in human myotubes following TMEM9B-AS1 silencing (Fig. 4d), suggesting that TMEM9B-AS1 could regulate MYC mRNA stability via a physical interaction with RNA-binding IGF2BP proteins. To probe this possibility, we performed an RNA pull-down experiment using the MYC-CRD region as bait to determine whether TMEM9B-AS1 is required for the interaction between IGF2BP1 or IGF2BP2 and the CRD region on MYC mRNA (illustrated in Fig 4e). Pull-down experiments were performed using cell lysates from human myotubes in the presence or absence of TMEM9B-AS1 silencing. The ability of MYC-CRD region to pull down IGF2BP1 and/or IGF2BP2 was assessed using Western blot analysis (Fig. 4f-j). Pulldown efficiencies and specificities for IGF2BP1 (Fig. 4g) and IGF2BP2 (Fig. 4h) were similar. Intriguingly, we observed a decrease in the pull-down efficiency of the MYC-CRD region in cells lacking TMEM9B-AS1 for IGF2BP1 (Fig. 4f, i), but not IGF2BP2 (Fig. 4f, j).

IGF2BP1 and IGF2BP2 are expressed in human myotubes and siRNA silencing of TMEM9B-AS1 decreases IGF2BP1 protein levels (Extended Data Fig. 5a-b), which indicates that TMEM9B-AS1 could also regulate the protein stability of IGF2BP1. We further investigated whether translation of MYC is altered in human myotubes in the absence of either TMEM9B-AS1, IGF2BP1, or IGF2BP2, but did not observe any changes in MYC protein levels (Extended Data Fig. 5c-d). IGF2BPs/IMPs are oncofetal proteins highly expressed in early development and in carcinogenesis^27^. IGF2BP2 has previously been implicated in the regulation of translation and ribosomal biogenesis ^28,29^. However, to our knowledge, here we provide the first data for the involvement of IGF2BP1 in ribosomal biogenesis. Common genetic variants on IGF2BP2 have been linked to type 2 diabetes and other metabolic phenotypes ^29,30^. Using the Type 2 Diabetes Knowledge Portal, ^31^ we observed that IGF2BP1 variants were strongly associated weight and moderately associated with height, glycated haemoglobin, and type 2 diabetes, providing new evidence for genetic association of IGF2BP1 and type 2 diabetes.

### IGF2BP1, but not IGF2BP2, regulates gene expression important for ribosomal biogenesis

We used the FANTOM 5 database to confirm the pattern and level of IGF2BP1 gene expression. Although transcript abundance is low, IGF2BP1 is expressed in several adult tissues including human skeletal muscle (Fig. 5a). We explored this finding further by dissecting data from the GTEX Portal ^32^ for single tissue/cell expression of IGF2BP1, TMEM9B-AS1 and MYC, which confirmed expression in skeletal muscle tissue, mostly within myocytes (Extended Data Fig. 6a-c). We validated this finding using another platform for single cell/nuclei data, CZ CELLxGENE Discover ^33^ and confirmed that IGF2BP1, TMEM9B-AS1 and MYC are expressed in skeletal muscle fibres and that TMEM9B-AS1 and IGF2BP1 have a similar expression pattern (Extended Data Fig. 6d). Moreover, we observed a higher expression level of IGF2BP1 in human myotubes compared to bulk skeletal muscle tissue from humans or rodents, as well as cultured rodent skeletal muscle cells (Extended Data Fig. 6e), suggesting possible species-specific roles for IGF2BP1 in human skeletal muscle.

**Figure 5.**
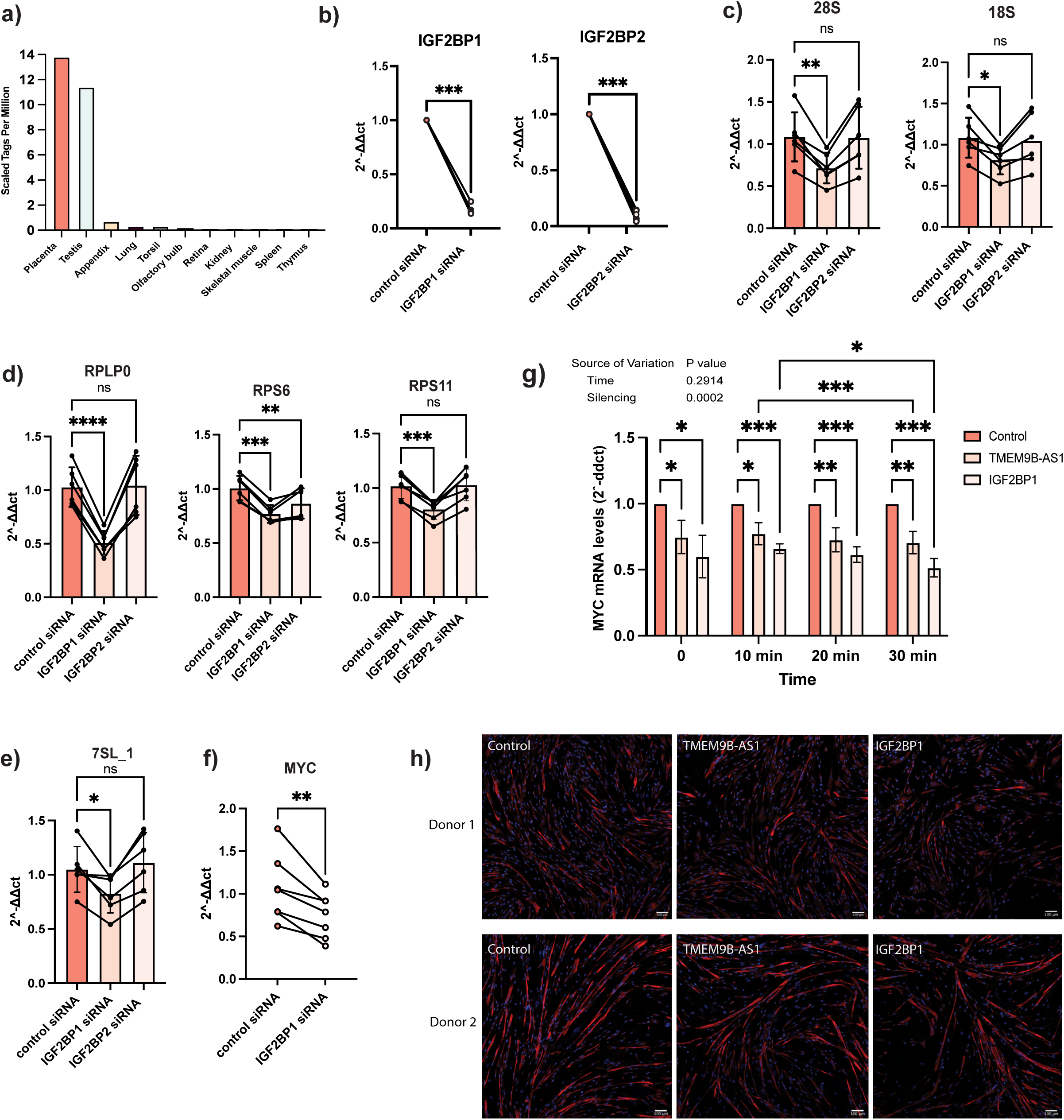
IGF2BP1 not IGF2BP2 regulates ribosomal biogenesis in human skeletal muscle cells. **a.** Expression of IGF2BP1 in different human tissues from FANTOM 5 data. **b.** siRNA silencing efficiencies of IGF2BP1 and IGF2BP2. QPCR results showing the expression levels of **c.** POLI, **d.** POLII and **e.** POLIII dependent ribosomal RNA levels upon IGF2BP1 and IGF2BP2 silencing. **a. f.** mRNA levels of MYC upon IGF2BP1 silencing. **g.** Results of the time course mRNA stability experiment of human myotubes treated with Actinomycin D for blocking transcription upon siRNA silencing of TMEM9B-AS1 and IGF2BP1**. QPCR** data normalized to the housekeeping gene(s) n=6, * p<0.05, ** p<0.01, *** p<0.001. **h.** Immunocytochemistry images showing Desmin protein levels (shown as red) in differentiated human myotubes in two different donors: Donor 1 and Donor 2. DAPI was used for the staining of the nuclei (shown as blue). Scale bar is 100 µm.

To further explore whether IGF2BPs are involved in the regulation of ribosomal biogenesis in human myotubes, we determined if siRNA silencing of IGF2BP1 or IGF2BP2 would phenocopy the TMEM9B-AS1-mediated reduction of genes important for ribosomal biogenesis (Figures 3b-d). Both IGF2BPs were efficiently silenced (Fig. 5b). However, the expression level of ribosomal RNAs/genes dependent on POL I (Fig. 5c), POL II (Fig. 5d) and POL III (Fig. 5e), were only decreased following silencing of IGF2BP1, apart from RPS6. These data are compatible with a model whereby TMEM9B-AS1 physically interacts with IGF2BP1, which in turn regulates mRNA stability of MYC by mediating the interaction of the MYC-CRD region and the RNA binding protein IGF2BP1. Indeed, IGF2BP1 regulates mRNA stability of MYC ^26,34^ in cell lines. Consistent with this, siRNA mediated silencing of IGF2BP1 reduced MYC mRNA (Fig. 5f). MYC mRNA is unstable with a short (15-30 min) half-life in human cells ^35,36^. Using this time window, we tested whether mRNA stability of MYC is affected by the silencing of TMEM9B-AS1 and IGF2BP1 upon blocking transcription using actinomycin D treatment. We observed a decline in MYC mRNA levels upon TMEM9B-AS1 and IGF2BP1 silencing between 10 min and 30 min after the treatment (Fig. 5g). Supporting our finding in Extended Data Fig. 2, we confirm with immunocytochemistry that silencing of TMEM9B-AS1 and IGF2BP1 reduces Desmin protein in human myotubes (Fig. 5h).

Our data provides evidence for a role of TMEM9B-AS1 in the regulation of ribosomal biogenesis and, in turn, protein synthesis. These processes are highly coordinated and essential for muscle mass and function, a feature that is impaired in type 2 diabetes ^6,37,38^. To validate whether reduced TMEM9B-AS1 would correspond to reduced ribosomal gene content we extracted all ribosomal protein coding genes present in our original RNA-SEQ data and compared expression in skeletal muscle from individuals with type 2 diabetes to individuals with normal glucose tolerance. Strikingly, most ribosomal genes were downregulated in skeletal muscle from individuals with type 2 diabetes (Fig. 6a), coincident with reduced TMEM9B-AS1. Since ribosomal biogenesis is essential for translational capacity and required for the maintenance of skeletal muscle mass, we further probed TMEM9B-AS1 expression in publicly available data sets. In older individuals predominantly performing either resistance or varying degrees of combined (i.e. resistance + endurance) exercise training, paradigms that result in ribosomal biogenesis and skeletal muscle hypertrophy, there was a non-significant tendency for increased TMEM9B-AS1 versus sedentary individuals (Fig. 6b, data dissected from GSE165630 ^39^). However, TMEM9B-AS1 expression was unchanged in a separate cohort of young resistance trained individuals compared to pre-training levels ^40^, measured across the initial phase of a training period characterized by marked accretion of ribosomal entities^40,41^ (Extended Data Fig. 7a). In contrast, sarcopenia represents a pathological condition of reduced skeletal muscle mass and analysis of a dataset including sarcopenia revealed downregulation of skeletal muscle TMEM9B-AS1 (Fig. 6c, data dissected from GSE11006 ^39^). Thus TMEM9B-AS1 appears to be involved in the maintenance of skeletal muscle mass, rather than regulating skeletal muscle hypertrophy.

**Figure 6.**
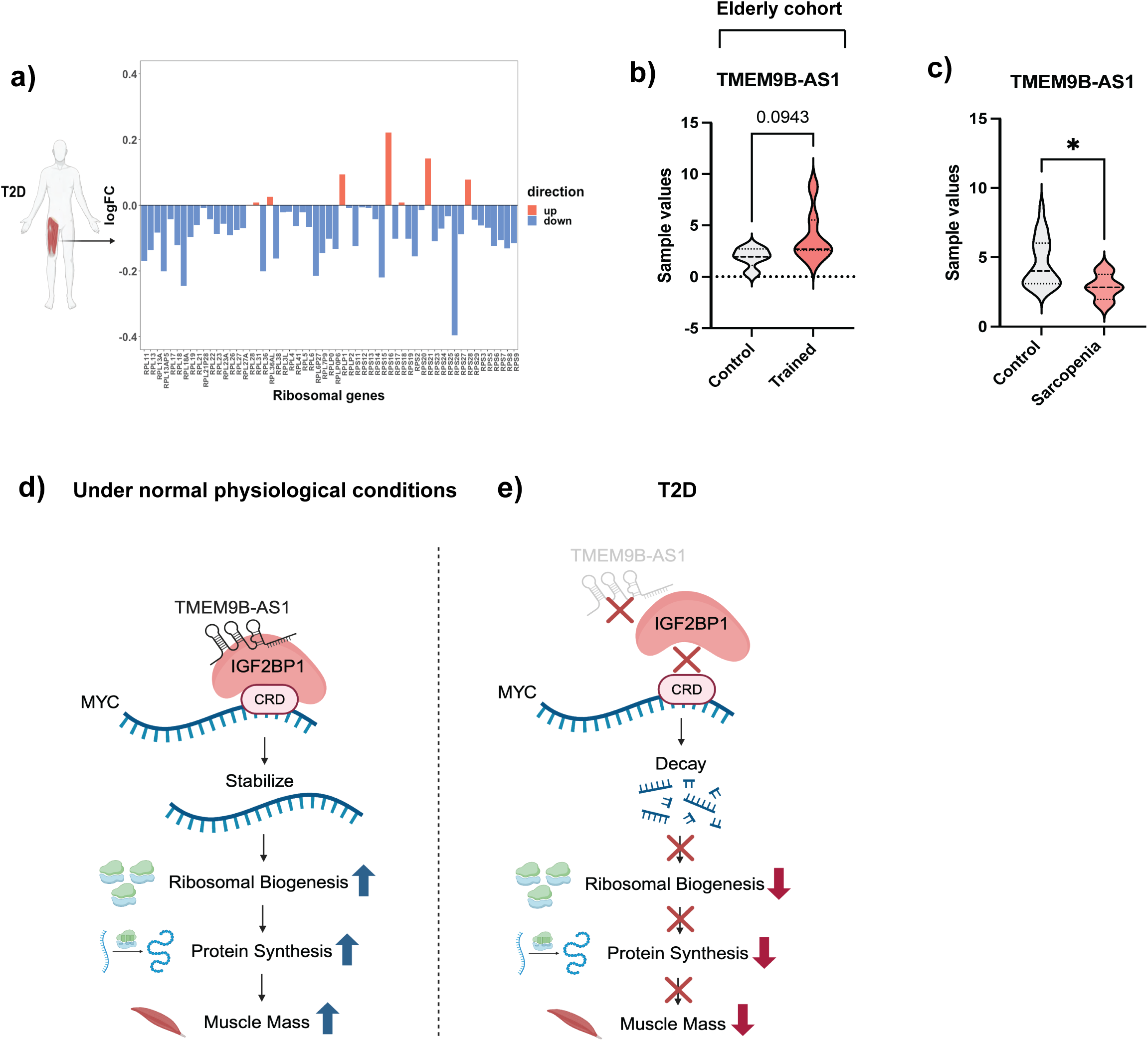
Supporting evidence for the regulation of ribosomal biogenesis and muscle mass by TMEM9B-AS1 in several human cohorts and the proposed model. **a.** Expression levels of ribosomal genes in the skeletal muscle of individuals with type 2 diabetes compared individuals with normal glucose tolerance. Data is extracted from the original RNA-SEQ data. **b.** The expression levels of TMEM9B-AS1 in GSE165630 dataset, derived from a clinical cohort following resistance or combined training (n=5 for control group and n=7 for trained group). **c.** TMEM9B-AS1 levels in people with sarcopenia compared to age-matched individuals without sarcopenia with normal muscle mass levels. GSE111006 dataset was used, and analysis were performed using GEO2R. Sample values are extracted and plotted. Welch two-tailed t test was used for the statistical analysis (n=32 for control group and n=4 for sarcopenia group). **d.** Under normal physiological conditions, TMEM9B-AS1 physically binds to the RNA-binding protein IGF2BP1 and mediates its interaction with MYC mRNA through its CRD region This facilitates the stability of MYC mRNA and eventually ribosomal biogenesis, as MYC is ultimately required for the expression of ribosomal RNAs/genes constituting ribosomal subunits. **e.** In type 2 diabetes, the decrease of TMEM9B-AS1 expression levels in the skeletal muscle disrupts the interaction between IGF2BP1 and MYC CRD is disrupted, which interferes with the mRNA stabilization of MYC mRNA. The subsequent decay eventually results in the decline of ribosomal biogenesis and in muscle mass.

### Proposed model

Based on our results we propose the following model (Fig. 6d-e): Under normal physiological conditions, TMEM9B-AS1 physically binds to the RNA-binding protein IGF2BP1. This binding facilitates the interaction between IGF2BP1 and MYC mRNA through the MYC-CRD region. This interaction, in turn, enhances the stability of MYC mRNA, which is essential for the transcription of ribosomal RNAs and genes that constitute ribosomal subunits (Fig. 6d). However, in the context of type 2 diabetes, and potentially other conditions associated with impaired skeletal muscle mass (e.g. sarcopenia), TMEM9B-AS1 expression is reduced. This reduction interferes with the interaction between IGF2BP1 and MYC-CRD, leading to a disruption in the mRNA stabilization process for MYC mRNA. Consequently, this disruption results in the decay of MYC mRNA and ultimately leads to a decline in ribosomal biogenesis, which impairs translational capacity and eventually the maintenance of muscle mass (Fig. 6e). The disrupted ribosomal biogenesis orchestrated by TMEM9B-AS1 holds physiological implications, potentially contributing towards the decrease in skeletal muscle mass often observed in individuals with type 2 diabetes. These findings open avenues for further research and therapeutic exploration in the context of type 2 diabetes and lncRNA function.

## Discussion

Here we provide a functional role for the lncRNA TMEM9B-AS1 in the regulation of ribosomal biogenesis, a crucial process for the maintenance and hypertrophy of skeletal muscle. While our analysis focuses on the role of TMEM9B-AS1 in skeletal muscle metabolism, this lncRNA may have a similar role in other tissues. Further investigation into additional protein interactors could deepen our understanding of TMEM9B-AS1’s coordination of ribosomal biogenesis and skeletal muscle integrity. To date, there are no marketed drugs directly targeting skeletal muscle mass. Thus, understanding the role of lncRNAs in the regulation of muscle mass could offer new insights into the preservation of skeletal muscle across the lifespan, and open novel therapeutic approaches for the management of type 2 diabetes and related metabolic disorders.

In addition to effects mediated by stabilizing MYC, TMEM9B-AS1 could facilitate other downstream processes via its regulation of IGF2BP1 protein stability. Moreover, our mass spectrometry data revealed several other protein interactors of TMEM9B-AS1 and further investigations are required to determine whether interaction of TMEM9B-AS1 with these proteins may be part of the mechanisms by which TMEM9B-AS1 modulates ribosomal biogenesis, protein synthesis, or other downstream effects. It could well be that the presence of TMEM9B-AS1 is necessary to sensitise muscle for growth such that when there is deficit of TMEM9B-AS1, it may be difficult to stabilise sufficient MYC mRNA required. In this scenario, the presence of sufficient TMEM9B-AS1 would be necessary for the muscle’s ability to respond to growth-stimulating stress.

### Limitations

Limitations of the present study include challenges in translating results to an in vivo model. Although we could overexpress TMEM9B-AS1 in skeletal muscle of mice to assess physiological outcomes on muscle mass and function, TMEM9B-AS1 is human specific - a feature shared with many lncRNAs ^42^. Despite mouse IGF2BP1 having 95% sequence similarity with the human homolog, RNA pulldown experiments using protein lysates from C2C12 mouse skeletal muscle cells using TMEM9B-AS1 as bait did not detect any interaction with mouse IGF2BP1. We also noted that expression levels of IGF2BP1 in mouse skeletal muscle cells was markedly reduced in comparison to human skeletal muscle cells, further underscoring species differences in this regulatory pathway. Thus, the TMEM9B-AS1/IGF2BP1 interaction appears specific for the regulation of ribosomal biogenesis in human skeletal muscle. In our clinical cohort, most participants with type 2 diabetes were medicated using Metformin and a substantial number were also using statins. Although no changes in TMEM9B-AS1 levels were noted following acute Simvastatin or Metformin treatment of human myotubes, we cannot exclude that chronic use of these drugs could lead to changes in vivo.

## Methods

### Human cohorts

Skeletal muscle samples from three previously published studies were used (^11^ ,^12,40^). All samples were collected with approval of the regional ethics boards and following permission of the participants.

### RNA sequencing analysis and differential expression

RNA sequencing data was used from our previous study ^11^ deposited in Gene Expression Omnibus Downstream analysis was conducted using R version v4.0. Data were filtered to keep only 13029 genes with >10 counts in at least 70% of samples from at least one group. The data set was then normalized using Trimmed Mean of M-Values (TMM) from the EdgeR package v3.32.1, and gene biotype annotation was collected from EnsDb.Hsapiens.v86 v2.99. Two samples were excluded from downstream analyses, as clustering of Manhattan distances between samples indicated that they were outliers. A linear model was fitted using limma v3.46. The linear model had one-way design as a combination of group (NGT or type 2 diabetes) and timepoint (Basal, Post-exercise, Rest), while subject pairing was accounted for by duplicate Correlation. Voom was applied to account for RNA sequencing heteroscedasticity. Obtained p values from contrasts of interest were adjusted by the Benjamini and Hochberg false discovery rate method.

### Growth and differentiation of primary human skeletal muscle cells

Primary cells were obtained from muscle biopsies taken from the vastus lateralis of healthy female and male volunteers ^43^ Myoblasts were allowed to proliferate in growth medium (F12/DMEM, 25 mM glucose, 20% FBS, 1% penicillin-streptomycin (Antibiotic-antimyotic, Thermo Fisher Scientific). To induce differentiation, cells were cultured in DMEM/M199 media supplemented with HEPES 20mM (Invitrogen), zinc sulfate (0.03 μg/mL), vitamin B12 (1.4 μg/mL; Sigma-Aldrich), insulin (10 μg/mL; Actrapid; Novo Nordisk), apo-transferrin (100 μg/mL; BBI Solutions), 0.5% FBS, and 1% penicillin-streptomycin (Anti-anti, Thermo Fisher Scientific). After 4-5 days, the cells were transitioned to post fusion medium, which consisted of DMEM/M199 (4:1), HEPES, zinc sulfate, vitamin B12, and 0.5% FBS, along with 1% penicillin-streptomycin. Cells were incubated in 7.5% CO2, humidified incubators at 37 °C, and medium was changed every other day during growth and differentiation.

### siRNA silencing

siRNA transfection was performed using either control siRNA, Stealth Control #1 medium GC (12935300, Invitrogen) or TMEM9B-AS1 Stealth siRNA (HSS162366, Thermo Fisher Scientific), IGF2BP1 Stealth siRNA (HSS173800, Thermo Fisher Scientific), IGF2BP2 Stealth siRNA (HSS173802, Thermo Fisher Scientific) (10nM final concentration) 6 days after induction of differentiation. A second transfection was conducted 48 hours later and all cellular assays were performed 2 days after the second transfection. Transfections were performed for 5 hours in OptiMEM reduced serum media with Lipofectamine® RNAiMAX (Invitrogen). Two days after the second transfection (i.e. day 10 of differentiation), cells were harvested, or subjected to downstream experiments.

### RNA-FISH combined with immunocytochemistry

RNA-FISH combined with immunocytochemistry assays for TMEM9B-AS1 and Desmin were performed using the ViewRNA Cell Plus Assay Kit Assay Kit (Invitrogen) according to the manufacturer’s instructions. Briefly, primary human skeletal muscle cells were seeded and differentiated on laminin (Sigma Aldrich) coated 4-well Nunc Lab-Tek II Chamber Slides (Thermo Scientific). At day 8 of differentiation, human myotubes were washed with phosphate buffered saline (PBS) containing RNase inhibitor and fixed with Fixation/Permeabilization Solution that is provided in the kit for 30 minutes at room temperature. Desmin antibody (1:500), secondary anti-goat antibody, Alexa fluor 546 (1:1000), and a probe against TMEM9B-AS1 (VA4-3102848-VCP, Thermo Fisher) were used for detection. First, antibody staining was performed by incubating the cells for 1 hour with the primary DESMIN antibody, followed by washes with 1x PBS with RNase inhibitor and 1 hour incubation with the secondary Alexa fluor 546 antibody. The RNA probe against TMEM9B-AS1 was diluted 1:100 and subsequently hybridized for 2 h at 40 °C. After hybridization and amplification steps with pre-amplifier, amplifier, and linked labelled probe, the probes were detected using Alexa^®^ Fluor dyes, according to the manufacturer’s instructions. Cells were mounted using ProLong™ Gold Antifade Mountant with DAPI (Invitrogen). Images were obtained using a Leica TCS SP8 confocal microscope (Leica Microsystems) and 63x oil objectives. Images were analyzed and and adjusted for brightness and contrast using ImageJ/Fiji software^44^.

### RNA isolation and cDNA synthesis

For RNA extraction and purification, the Quick-RNA kit from Zymo Research was used. The cells used in experiments were washed with PBS, lysed, scraped into the lysis buffer provided in the kit and snap frozen with liquid nitrogen. To isolate RNA, the samples were allowed to thaw to room temperature before proceeding with kit purification and isolation. RNA concentration and purity were assessed using spectrophotometry (NanoDrop, Thermo Scientific). For cDNA synthesis the High Capacity cDNA Reverse Transcription kit (Thermo Fisher Scientific) was used according to manufacturer’s instructions, with 1μg of purified total RNA as starting material. In short, a combination of random primers, deoxynucleotide triphosphate, RT buffer, and the MultiScribe Reverse Transcriptase enzyme were added to the RNA samples. Reverse transcription was performed, commencing with a 10-minute step at 25°C, followed by a 120-minute incubation at 37°C to facilitate cDNA polymerization. Subsequently, the samples were subjected to a 5-minute heat treatment at 85°C to halt the reaction.

#### QPCR experiments

Gene expression was determined by real-time QPCR using the Viia7 system (Thermo Fisher Scientific). Results were then analyzed using the Relative gene expression was calculated using the ΔΔCt method according to the housekeeping/reference genes with stable expression levels. Unless otherwise specified, the geometric mean of two or more housekeeping genes was used for analysis. QPCR was performed using TaqMan Fast Universal PCR Master Mix (Applied Biosystems) or Fast SYBR™ Green Master Mix (Applied Biosystems) and primers or pre-designed probes (Thermo Fisher Scientific) are listed in Supplementary Table 1.

#### Western Blot experiments

Human skeletal muscle cells were deprived of serum for 4 hours, and subsequently, they were exposed to insulin for a 10-minute period. Cells were then lysed and homogenized using a lysis buffer consisting of 137 mM NaCl, 2.7 mM KCl, 1 mM MgCl2, 0.5 mM Na3VO4, 1% (vol/vol) Triton X-100, 10% (vol/vol) glycerol, 20 mM Tris, 1 mM EDTA, 1 mM PMSF, 1% (vol/vol) Protease Inhibitor Cocktail Set 1 (Merck Millipore) and 1x PhosSTOP (Roche). These lysates were gently rotated at 4 °C for 30 minutes before subjecting to centrifugation at 12,000 × g at 4 °C for 15 minutes. The protein concentration was measured using the Pierce BCA Protein Assay Kit (Thermo Fisher Scientific), and equal quantities of protein were mixed with Laemmli buffer or 5x Pierce Lane Marker Reducing Sample Buffer (Thermo Fisher Scientific). The samples were boiled for 5-10 minutes and then separated using SDS-PAGE with 4-12% Criterion XT Bis-Tris Gels (Bio-Rad). The proteins were transferred to PVDF membranes (Merck Millipore), and Ponceau S (Sigma-Aldrich) staining was performed. Following this, the membranes were blocked in 10% milk in TBS-T (10 mM Tris-HCl, 100 mM NaCl, 0.02% Tween 20) for 1 hour at room temperature and subsequently incubated at 4 °C overnight with 1:1000 diluted primary antibodies (listed in Supplementary Table 1). The membranes were then washed with TBS-T, exposed to suitable secondary antibodies in 5% milk containing TBS-T with a dilution of 1:10000 for 1 hour at room temperature, and then washed again with TBS-T. Chemiluminescence Detection was conducted using Amersham ECL or Amersham ECL select Western Blotting Detection Reagent (GE Healthcare). Protein content was quantified through densitometry (utilizing QuantityOne software from Bio-Rad or Image J). The complete scanned blots are available in the Extended Data Fig. 8-9.

### SunSET Assay

The protocol for SunSET assay is adapted from Schmidt et al. 2009 ^45^. Human myotubes were washed once with PBS and starved in low glucose (1g/L) DMEM for 4-5 hours. Puromycin (1uM final) was then added to the media and incubated for 30 minutes. Prior to the harvest, cells were washed with ice cold PBS and scraped into lysis buffer consisting of 137 mM NaCl, 2.7 mM KCl, 1 mM MgCl2, 1% (vol/vol) Triton X-100, 10% (vol/vol) glycerol, 20 mM Tris, 1 mM EDTA, 1 mM PMSF, 1% (vol/vol) Protease Inhibitor Cocktail Set 1 (Merck Millipore) and 1x PhosSTOP (Roche). These lysates were gently rotated at 4 °C for 30 minutes before being subjected to centrifugation at 12,000 × g at 4 °C for 15 minutes. Supernatant was collected and stored at -80 °C until sample preparation for immunoblot analysis.

#### In vitro transcription and 3’ End Desthiobiotinylation

gBlocks gene fragments (Integrated DNA Technologies) were ordered for TMEM9B-AS1 and MYC-CRD that contains the sequence of T7 RNA polymerase promoter region for in vitro transcription. They were prepared and kept according to the manufacturer’s instructions.

T7 RNA polymerase (New England Biolabs) and components for the standard RNA synthesis were mixed and incubated at 37°C for 3-4 hours. Immediately after this incubation, 0.1u/µl Turbo™ DNase (Invitrogen) was added and the mixture was incubated for an additional 15 minutes at 37°C. Next, 25 µl of RNase free water was added to the mixture and transcribed RNA was eluted using the RNA Clean & Concentrator kit (Zymo Research). Samples were assessed for their length and concentration by Bioanalyzer (Agilent) and Nanodrop (Thermo Scientific).

Pierce RNA 3’ End Desthiobiotinylation Kit (Thermo Scientific) was used for the biotinylation of the transcribed RNA for the pulldown experiments. Approximately 30pmol of RNA was used for desthiobiotinylation. To relax the secondary structure, RNA was heated for 3-5 minuntes at 85°C in the presence of 25% DMSO (provided in the kit) and cooled down rapidly on ice. Ligation reactions for desthiobiotinylation were prepared according to the manufacturer’s instructions and were incubated overnight at 16°C. The next day, desthiobitinylated RNA was extracted using chloroform:isoamyl alcohol and the RNA pellet was resuspended in nuclease-free water. The final concentration of desthiobiotinlylated RNA was measured using Nanodrop. 30 ρmol of desthiobiotinylated control or test RNA was used for purifications.

#### RNA pulldown experiments

15 cm dishes were seeded with primary human skeletal muscle cells and differentiation was done in these plates. Cells were harvested at day 9 after differentiation for the pulldown experiments. For the siRNA silencing experiments cell media were changed to post fusion at day 5. And, double transfections were performed at day 6 and day 8 of differentiation. Cells were transfected with control stealth RNAi (Control #1, Medium GC, Invitrogen) or with TMEM9B-AS1 stealth RNAi (HSS162366, Thermo Fisher Scientific). Then, 10 days after the start of differentiation, cells were scraped and harvested in 300 ul IP lysis buffer (25 mM Tris-HCl pH 7.4, 150 mM NaCl, 1% NP-40, 1 mM EDTA, 5% glycerol), supplemented with Protease Inhibitor Cocktail Set 1 (Merck Millipore) and RNase inhibitors (Applied Biosystems). Cell lysates were gently rotated at 4 °C for 30 minutes before being subjected to centrifugation at 12,000 × g at 4 °C for 15 minutes and supernatant was kept on ice for the pulldown experiments. 30 µL of the lysate was used per pulldown condition. Proteins were pulled down either on TMEM9B-AS1-bound Streptavidin beads or control RNA (PolyA)-bound Streptavidin beads. Pierce Magnetic RNA-Protein Pull-Down Kit was used for the experiments (Thermo Scientific). For the rest of the protocol manufacturer’s instructions and recommendations were followed.

### Sample preparation for Mass Spectrometry

Protein eluents (ca 50 µL) were supplemented with 1 µL of 1M ammonium bicarbonate (AmBic) and digested with 2 µL of 0.05 µg/µL sequencing grade trypsin (Promega) in 100 mM AmBic, incubating at 37°C overnight with shaking at 450 rpm on a thermal heater. The digestion was stopped with 1 µL of cc. formic acid (FA), then the samples were transferred to a vial and dried in a vacuum concentrator (Eppendorf).

### Liquid Chromatography-Tandem Mass Spectrometry Data Acquisition

Peptides were reconstituted in 7 µL of solvent A (0.1% FA in water) and 5 µL was injected on a 50 cm long EASY-Spray C18 column (Thermo Fisher Scientific) connected to an UltiMate 3000 nanoUPLC system (Thermo Fisher Scientific) using a 60 min long gradient: 4-26% of solvent B (98% acetonitrile, 0.1% FA) in 60 min, 26-95% in 5 min, and 95% of solvent B for 5 min at a flow rate of 300 nL/min. Mass spectra were acquired on a Q Exactive HF hybrid quadrupole Orbitrap mass spectrometer (Thermo Fisher Scientific) ranging from m/z 375 to 1500 at a resolution of R=120,000 (at m/z 200) targeting 5×106 ions for maximum injection time of 100 ms, followed by data-dependent higher-energy collisional dissociation (HCD) fragmentations of precursor ions with a charge state 2+ to 7+, using 30 s dynamic exclusion. The tandem mass spectra of the top 17 precursor ions were acquired with a resolution of R=30,000, targeting 2×105 ions for maximum injection time of 54 ms, setting quadrupole isolation width to 1.4 Th and normalized collision energy to 28%.

### Protein identification

Acquired raw data files were analysed using Proteome Discoverer v2.4 (Thermo Fisher Scientific) with Mascot Server v5.1 (MatrixScience, UK) against human protein database (SwissProt). A maximum of two missed cleavage sites were allowed for full tryptic digestion, while setting the precursor and the fragment ion mass tolerance to 10 ppm and 0.02, respectively. Carbamidomethylation of cysteine was specified as a fixed modification. Oxidation on methionine, deamidation of asparagine and glutamine were set as dynamic modifications.

### Selection of the protein interactors of TMEM9B-AS1

Proteomics data of control and TMEM9B-AS1 pulldowns were compared. First proteins that only appeared in TMEM9B-AS1 pulldown were identified for each experiment/donor. Then, peak intensities for the peptides were used to identify proteins having abundance at least 10-fold in TMEM9B-AS1 pulldown in comparison to the control pulldown for each experiment/donor. Combined lists of protein interactors were created for each donor (three donors in total) and these lists were compared with each other and overlapping proteins were determined as specific interactors as it was shown venn diagrams in figure 3e.

### Glucose oxidation assay

Differentiated human myotubes were serum-starved for 2 hours in low-glucose DMEM (Gibco, 5.5mM) Then, the media was changed to the one supplemented with D-[U-^14^C]glucose in the presence or absence of 1 µM FCCP (Sigma-Aldrich). An empty cup was put in each well and plates were sealed and incubated for 4 hours at 37°C in 0% CO_2_ incubator. After the incubation, media was acidified (1:8 vol 2 M HCl) and the liberated ^14^CO_2_ was collected for an hour in a well containing 2 M NaOH. The quantification for the liberated ^14^CO_2_ was determined by scintillation counting. Cells were harvested in 400 µl 0.5 M NaOH, pH was neutralized by the addition of 100µl of 2 M HCl. Protein content was determined by BCA Protein Assay Kit (Thermo Fisher Scientific).

### Fatty acid (palmitate) oxidation assay

Differentiated human myotubes were serum-starved for 2 hours in low-glucose DMEM (Gibco, 5.5mM) and then were incubated for 6 hours in low-glucose DMEM supplemented with 25μM of palmitate, including 0.078μM [9,10-^3^H(N)] palmitate (PerkinElmer) and 0.04% BSA and in the presence or absence of 1 µM FCCP (Sigma-Aldrich). Supernatant was collected and incubated with 800µl of 10% activated charcoal in 20mM Tris-HCl buffer, pH 7.5 for 30 minutes. Samples were centrifuged at 13,000g for 15 minutes. 200µl of the supernatant was counted in a liquid scintillation counter. Protein content of the cells was measured using the BCA Assay Kit (Thermo Fisher Scientific) and the counts were normalized to the protein content of the cells.

### Glucose uptake assay

Differentiated human myotubes were serum-starved for 4 hours in low-glucose (Gibco, 5.5 mM) and then were incubated for 60 minutes, at 37°C in the presence or absence of 100nmol/l insulin. Then, cells were treated with glucose- and serum-free DMEM (Gibco) with the addition of 2-[1,2-3H]deoxy-D-glucose (Moravek) and 10μmol/L unlabeled 2-deoxy-d-glucose for 15 minutes. Upon incubation, cells were washed three times with ice-cold PBS and were lysed in 1 ml 0.03% SDS. 500µl of this lysate was transferred to the scintillation vials and counted in a 1414 WinSpectral Liquid scintillation counter. Protein content of the cells was measured using the BCA Assay Kit (Thermo Fisher Scientific) and the counts were normalized to the protein content of the cells.

### mRNA stability assay

Upon siRNA transfections at day 6 and 8, cells were treated either with Actinomycin D (Sigma) (4nM final concentration) or vehicle. Samples were collected at basal, 10, 20 and 30 minutes after the treatment in lysis buffer for RNA isolations and were immediately frozen in liquid nitrogen. EZNA total RNA kit (Omega) was used for RNA isolation and High-Capacity cDNA Reverse Transcription kit (Thermo Fisher Scientific) was used for cDNA synthesis by following the manufacturer’s instructions. MYC mRNA levels were measured by QPCR and data was normalized to the expression levels of the housekeeping gene, TBP.

### Immunocytochemistry

Primary human skeletal muscle cells were seeded and differentiated on 4-well Nunc Lab-Tek II Chamber Slides (Thermo Scientific) coated with laminin (Sigma Aldrich) according to manufacturer’s instructions. Double transfection was performed for siRNA silencing of TMEM9B-AS1 and IGF2BP1 at day 6 and day 8 after the differentiation has started. At day 10, cells were fixed for 10 minutes with 4% paraformaldehyde, quenched with 0.1 M glycine for 10 minutes at room temperature. Cells were washed 2 times with PBS and were permeabilised for 3 minutes with 0.1% Triton X-100 in PBS. After 2 washes with PBS, samples were blocked for 30 minutes in 1% BSA in PBS and incubated with primary antibody for Desmin (1:500 diluted in PBS) overnight at 4°C. Upon 2 washes with 0.025% Tween 20 in PBS secondary antibody incubation was performed for 90 minutes at room temperature using Alexa fluor 546 (1:1000 diluted in PBS). After 2 washes with 0.025% Tween 20 in PBS, cells were mounted using ProLong™ Gold Antifade Mountant with DAPI (Invitrogen). Images were obtained using a Leica Stellaris 5 LIA confocal microscope (Leica Microsystems) and 20x dry objectives. A random region was selected on the slides and 9 pictures were taken including and surrounding that area. Images were stitched together to have the final field of view for comparisons. Images were analyzed and adjusted for brightness and contrast using ImageJ/Fiji software ^44^.

### STRING, KEGG Pathway and GO term analysis for protein interactors

STRING database versions 11.5 and 12.0 were used for the protein-protein interaction networks and KEGG Pathway and GO term enrichment analysis ^46^. Default settings were used for the predictions with the usage of confidence as meaning of network edges. The line thickness represents the strength of the interaction.

#### GTEx expression overview

Bulk tissue gene expression information for TMEM9B-AS1was achieved from the GTEx portal^47^. Data was extracted for the sex specific expression of TMEM9B-AS1 in different metabolic tissues.

#### FANTOM 5 expression overview

Expression of IGF2BP1 in different human tissues, data was derived from the FANTOM5 dataset^48^ that was represented on the Human Protein Atlas version 23.0 ^49^.

#### Gene expression profiles for TMEM9B-AS1 from Gene Expression Omnibus (GEO)

GEO2R with the default settings was used to analyse the datasets GSE11006 and GSE165630. Specific gene expression profiles for TMEM9B-AS1 were dissected and sample values were extracted. Statistical analyses were conducted using Welch t-test.

### Statistical analysis

GraphPad Prism 10 software (GraphPad Software Inc., CA) was used for the statistical analyses. Normal distribution of the data was tested with Shapiro–Wilk test. For the data sets that were normally distributed, differences between two groups were assessed using paired two-tailed t-tests. For the datasets that are not following normal distribution, Mann Whitney test or Welch t-test were applied to assess differences between two groups. Differences between more than two groups were assessed by using one-way analysis of variance (ANOVA) followed by Dunnett’s test or Tukey’s test for post hoc pair-wise comparisons. Individual data points are shown in the plots and graphs and the variation is presented as SD together with the mean of the data sets. P values for significances are as follows: *p < 0.05; **p ≤ 0.01; ***p ≤ 0.001; ****p ≤ 0.0001.

### Data availability

The mass spectrometry proteomics data have been deposited to the ProteomeXchange Consortium via the PRIDE ^50^ partner repository with the dataset identifier PXD053490.

## Acknowledgements

We thank Proteomics Biomedicum at Karolinska Institute for mass spectrometry analyses, and the Biomedicum Imaging Core Facility (BIC) and Jianping Liu from The Single Cell Core Facility for Flemingsberg campus (SICOF) at Karolinska Institute for the confocal imaging. We thank Emmanuelle Charrin at Karolinska Institute for the help with the RNA-FISH experiments and the final confocal images. We thank Bo Falck Hansen at Novo Nordisk for the scientific discussions. Finally, we thank all Zierath and Krook lab members for helpful discussions. Open access funding is provided by Karolinska Institute. Figures were generated using Biorender.

## Author information

### Contributions

I.S., J.R.Z., and A.K. conceived and designed the study; I.S., J.S., E.C and K.L., performed experiments; I.S. and M.S. analysed RNA sequencing data; S.E supervised K.L and designed experiment, I.S. analysed data and I.S. and A.K. interpreted results of experiments; I.S. prepared figures and tables, and drafted the manuscript; I.S., J.R.Z., and A.K. finalized the manuscript. All authors read and approved the manuscript.

### Ethical declarations

All human samples collected with approval of the regional ethics boards and following permission of the participants

### Grants

Ilke Sen was supported by a Novo Nordisk postdoctoral fellowship run in partnership with Karolinska Institute. This study was supported by grants from EFSD/Lilly Young Investigator Award, the Swedish Diabetes Foundation and the Swedish Research Council and Region Stockholm.

## Extended Data Figures

**Extended Data Fig. 1.**
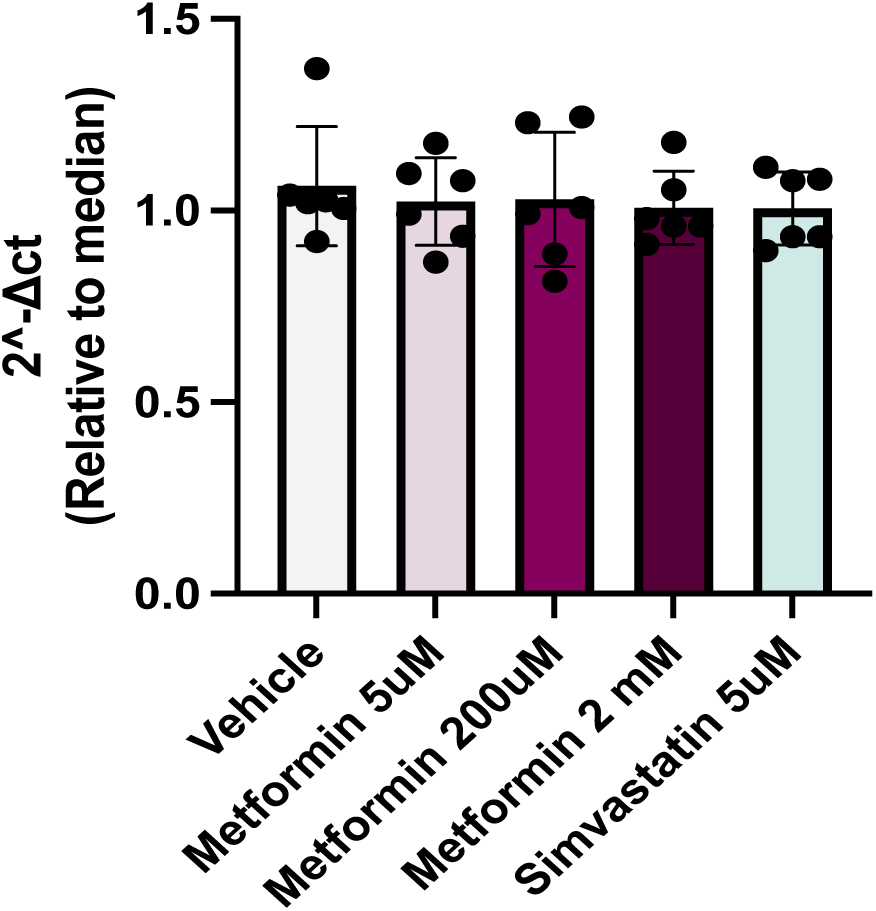
TMEM9B-AS1 levels do not change in human skeletal muscle cells upon Metformin and Simvastatin treatment. QPCR results showing the expression levels of TMEM9B-AS1 upon treatment of human myotubes with Metformin at three different doses; low (5uM), medium (200uM) and high (2mM) and Simvastatin (5uM). Data are normalized to the housekeeping/reference genes B2M and TBP n=6.

**Extended Data Fig. 2.**
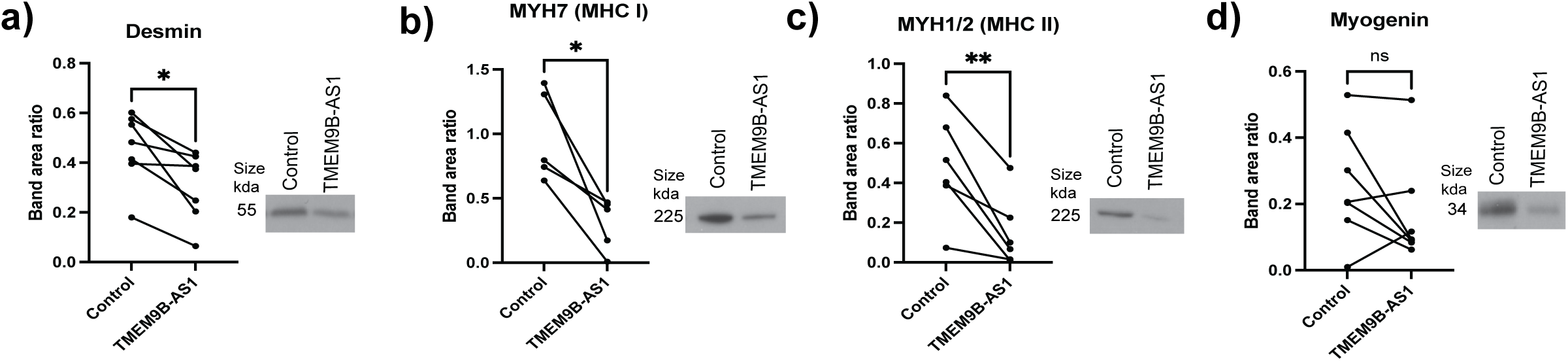
TMEM9B-AS1 regulates the expression of essential proteins that are required for muscle differentiation and structure. **a.** Western blot results of Desmin (n=7), **b.** MYH7 (MHC I) (n=5), **c.** MYH1/2 (MHC II) (n=6), **d.** Myogenin (n=7) in control and TMEM9B-AS1 siRNA treated human myotubes. Band area ratios were calculated by the normalization of the band intensity of the protein of interest to a selected ponceau band. *p<0.05, **p<0.01.

**Extended Data Fig. 3.**
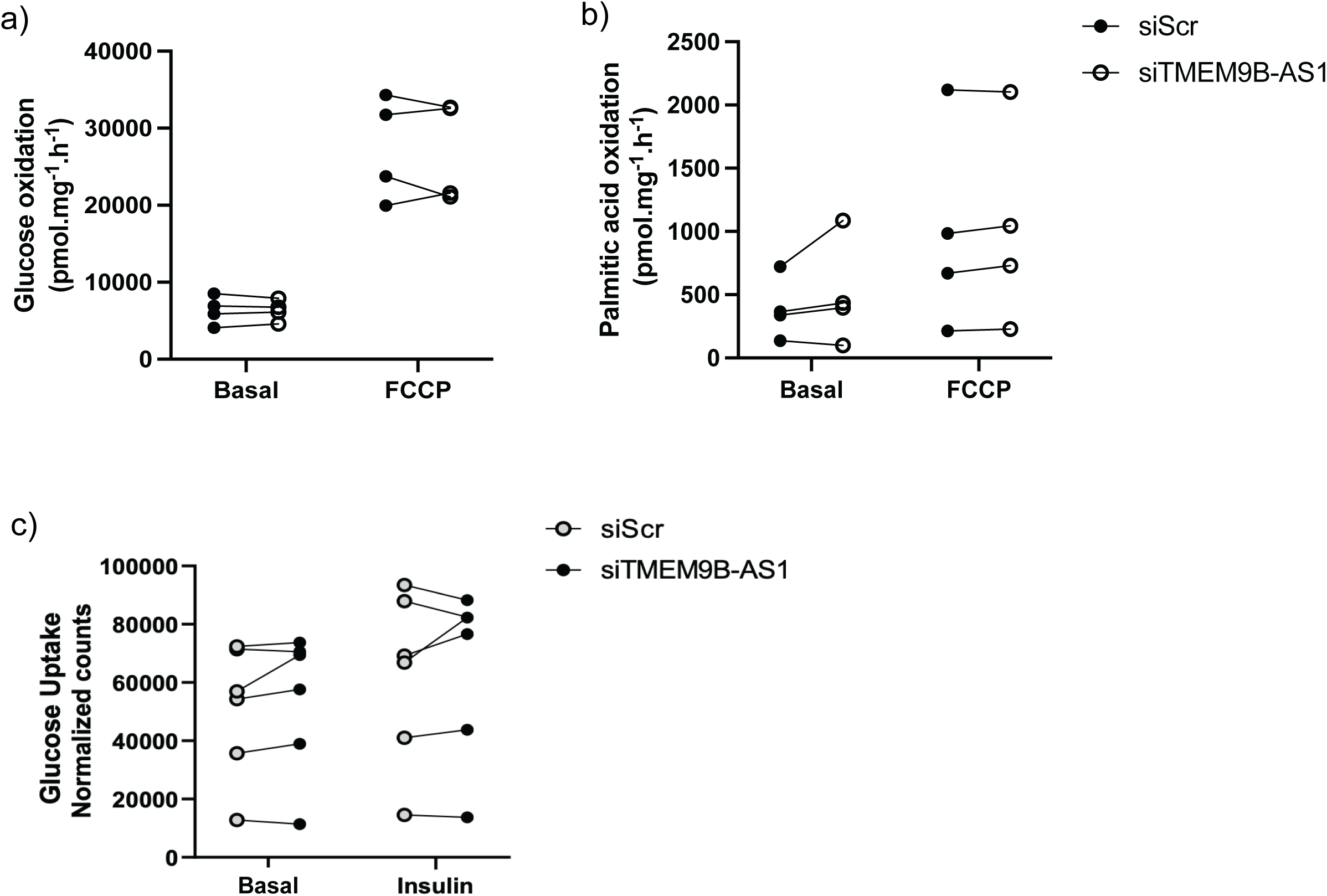
TMEM9B-AS1 does not regulate glucose or lipid metabolism in human myotubes. **a.** Glucose oxidation (n=4), **b.** Palmitic acid (lipid) oxidation (n=4) and **c.** Glucose uptake (n=6) assay results in human myotubes. Measeurements were done at baseline or upon a stimulation with FCCP (for oxidation assays) and insulin (for glucose uptake). Results were normalized to total protein content of the myotubes.

**Extended Data Fig. 4.**
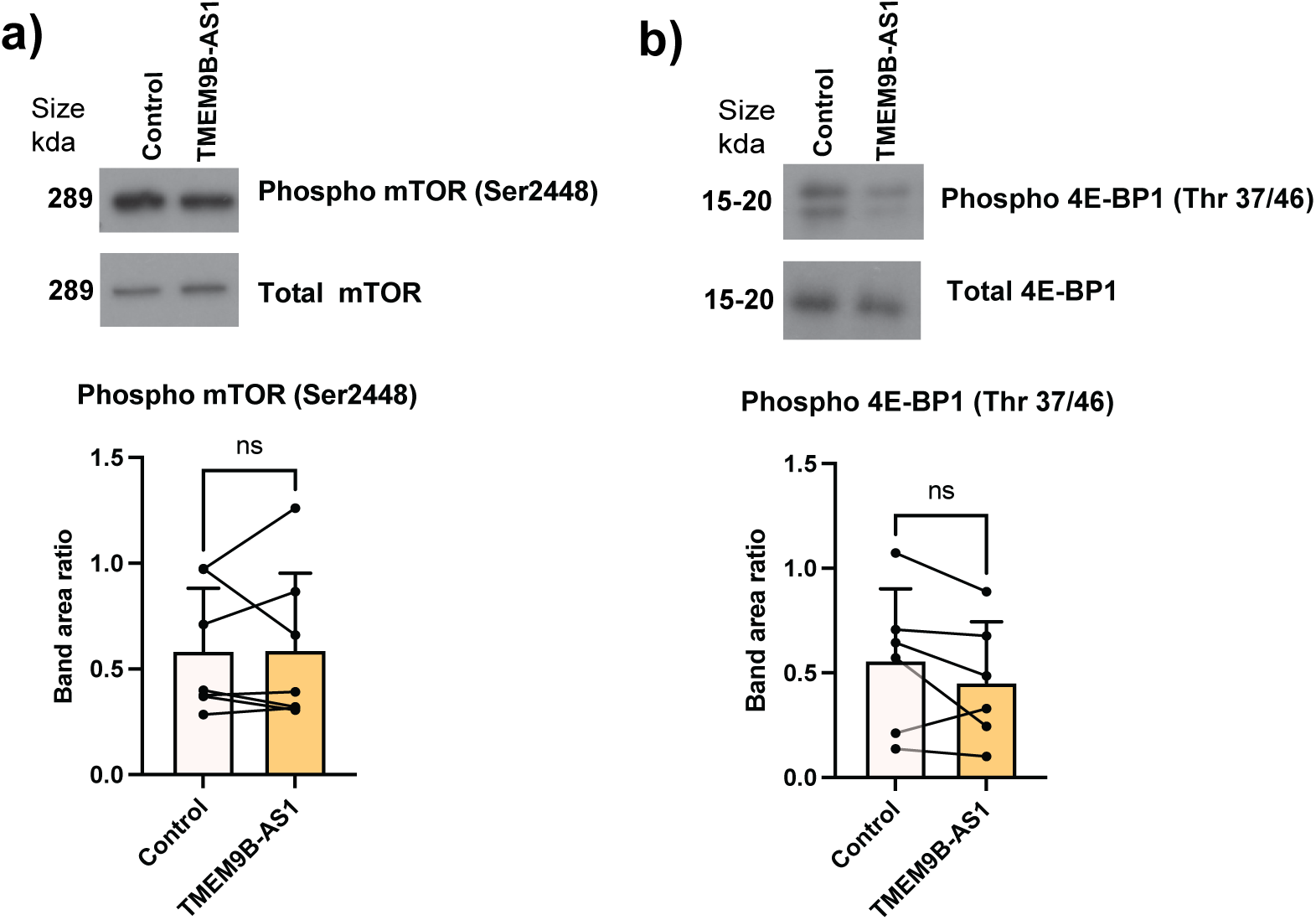
TMEM9B-AS1 silencing does not change the phosphorylation level of mTOR nor 4E-BP1 in human myotubes. Western blot results showing the phosphorylation level of **a.** mTOR (n=6), **b.** 4E-BP1 (n=7) upon TMEM9B-AS1 silencing. Band intensities were normalized to a Ponceau band. Then, phosphorylation levels were normalized to total protein levels.

**Extended Data Fig. 5.**
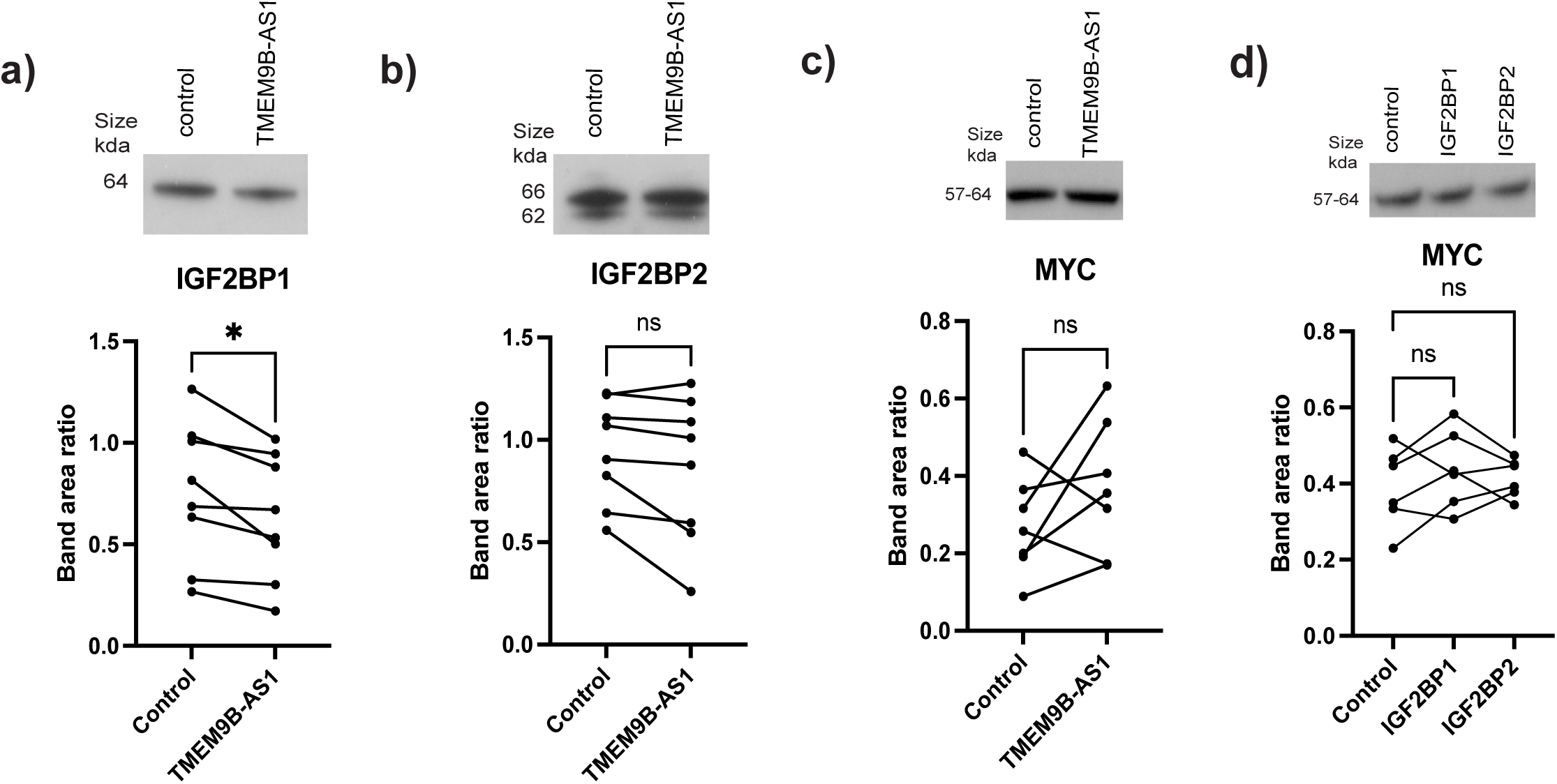
IGF2BP1 protein level is regulated by TMEM9B-AS1 and MYC translation is not affected by the loss of TMEM9B-AS1 nor IGF2BPs. Western blot results showing the protein levels of **a.** IGF2BP1(n=8) **b.** IGF2BP2 (n=8) **c.** MYC (n=6) upon TMEM9B-AS1 silencing in human myotubes. **d.** Western blot results for MYC protein levels upon IGF2BP1 and IGF2BP2 siRNA silencing (n=6). Band area ratios were calculated by the normalization of the band intensity of the protein of interest to a selected ponceau band. *p<0.05.

**Extended Data Fig. 6.**
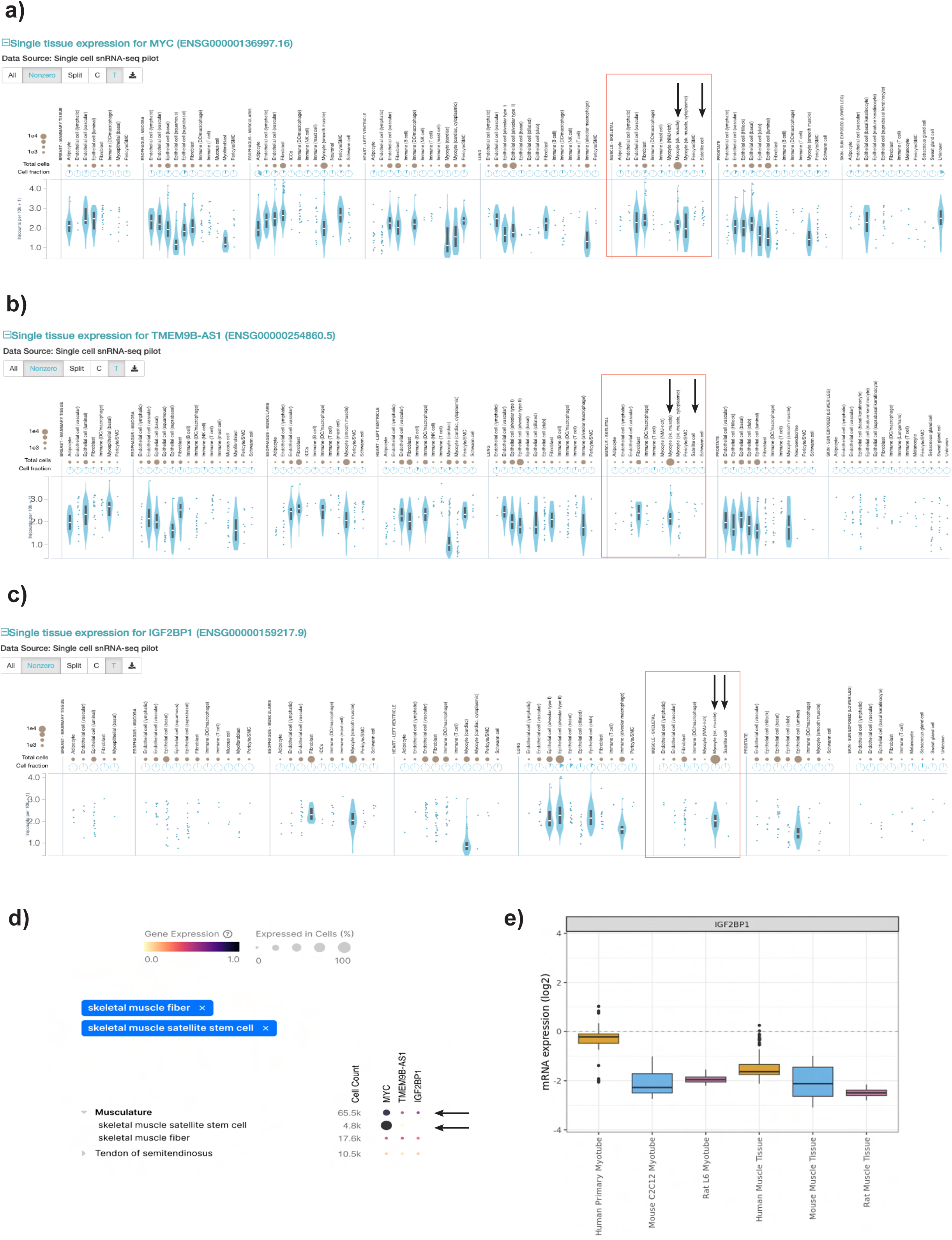
IGF2BP1 is expressed in human skeletal muscle cells/fibers and its expression level is the highest in human myotubes compared to the rodent skeletal muscle cells and tissues. Single tissue expression pattern of **a.** MYC, **b.** TMEM9B-AS1, and **c.** IGF2BP1 on GTEX database ^32^ Arrows are indicating the expression level of TMEM9B-AS1, IGF2BP1 and IGF2BP2 in myocytes and satellite cells in skeletal muscle tissue. **d.** Expression level of MYC, TMEM9B-AS1 and IGF2BP1 in skeletal muscle fibers and satellite stem cells on the single-cell data platform, CZ CELLxGENE Discover ^33^. **e.** Comparison of the expression level of IGF2BP1 mRNA in human and rodent skeletal muscle cells and tissues.

**Extended Data Fig. 7.**
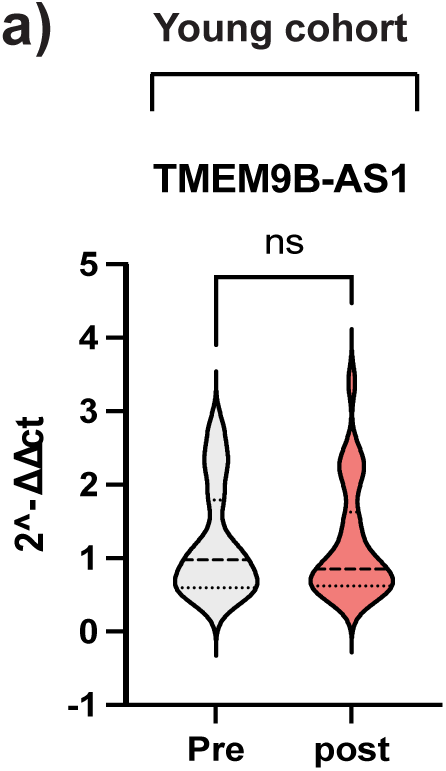
TMEM9B-AS1 levels do not change in human skeletal muscle of young individuals upon resistance training. **a.** QPCR results of TMEM9B-AS1 in skeletal muscle of young individuals before and after a resistance training intervention (n=26 for pre-training group and n=26 for post-training group).

**Extended Data Fig. 8.**
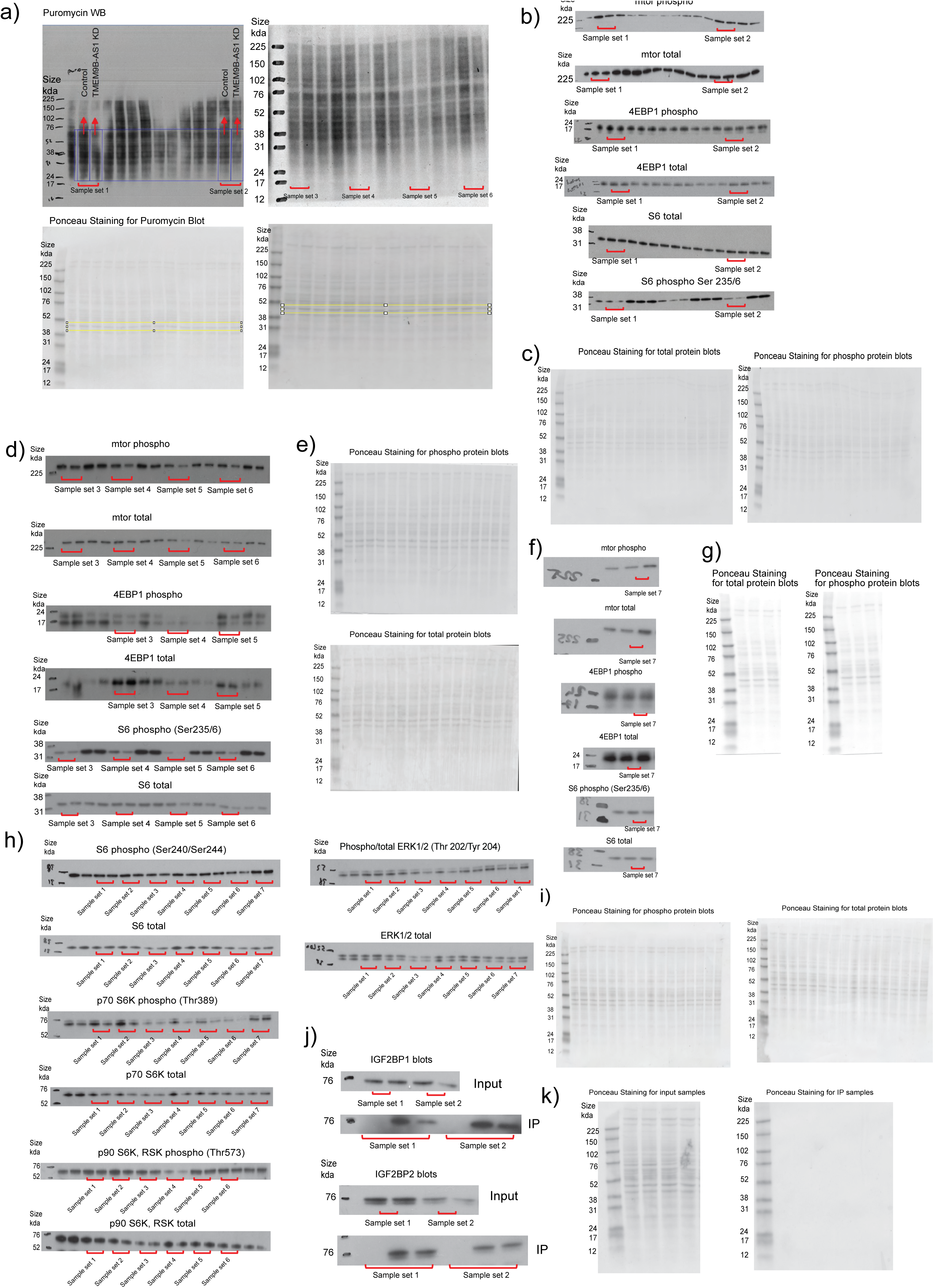
All membranes part I. Uncut membranes and ponceau staining with the information about the sample sets/replicates for **a.** Puromycin western blot experiment in figure 1f, **b.** Uncut membranes for sample set 1 and 2 for phosphorylated and total mTOR, 4EBP1 (Extended Data Fig. 4a-b), S6 protein (for Ser235/236 in figure 2g), **c.** Ponceau staining for sample set 1 and 2 for Phosphorylated and total mTOR, 4EBP1 (Extended Data Fig. 4a-b), S6 protein (for Ser235/236 in figure 2g). **d.** Uncut membranes for the. sample sets 3-6 for phosphorylated and total mTOR, 4EBP1 (Extended Data Fig. 4a-b), S6 protein (for Ser235/236 in figure 2g), **e.** Ponceau staining for sample the sets 3-6 phosphorylated and total mTOR, 4EBP1 (Extended Data Fig. 4a-b), and S6 protein (for Ser235/236 in figure 2g). **f.** Uncut membranes for the. sample sets 7 for phosphorylated and total mTOR, 4EBP1 (Extended Data Fig. 4a-b), and S6 protein (for Ser235/236 in figure 2g), **g.** Ponceau staining for sample the set 1-7 for phosphorylated and total mTOR, 4EBP1 (Extended Data Fig. 4a-b), and S6 protein (for Ser235/236 in figure 2g). **h.** Uncut membranes for the sample sets 1-7 for phosphorylated and total S6 protein (for Ser240/244 in figure 2h), p70 S6K (Fig. 2i), p90 S6K, RSK (Fig. 2j) and ERK1/2 (Fig. 2k) **i.** Ponceau staining for the sample sets 1-7 for phosphorylated and total S6 protein (for Ser240/244 in figure 2h), p70 S6K (Fig. 2i), p90 S6K, RSK (Fig. 2j) and ERK1/2 (Fig. 2k). **j.** Uncut membranes for the RNA pulldown experiments in figure 4f. **k.** Ponceau staining for the RNA pulldown experiments in figure 4f.

**Extended Data Fig. 9.**
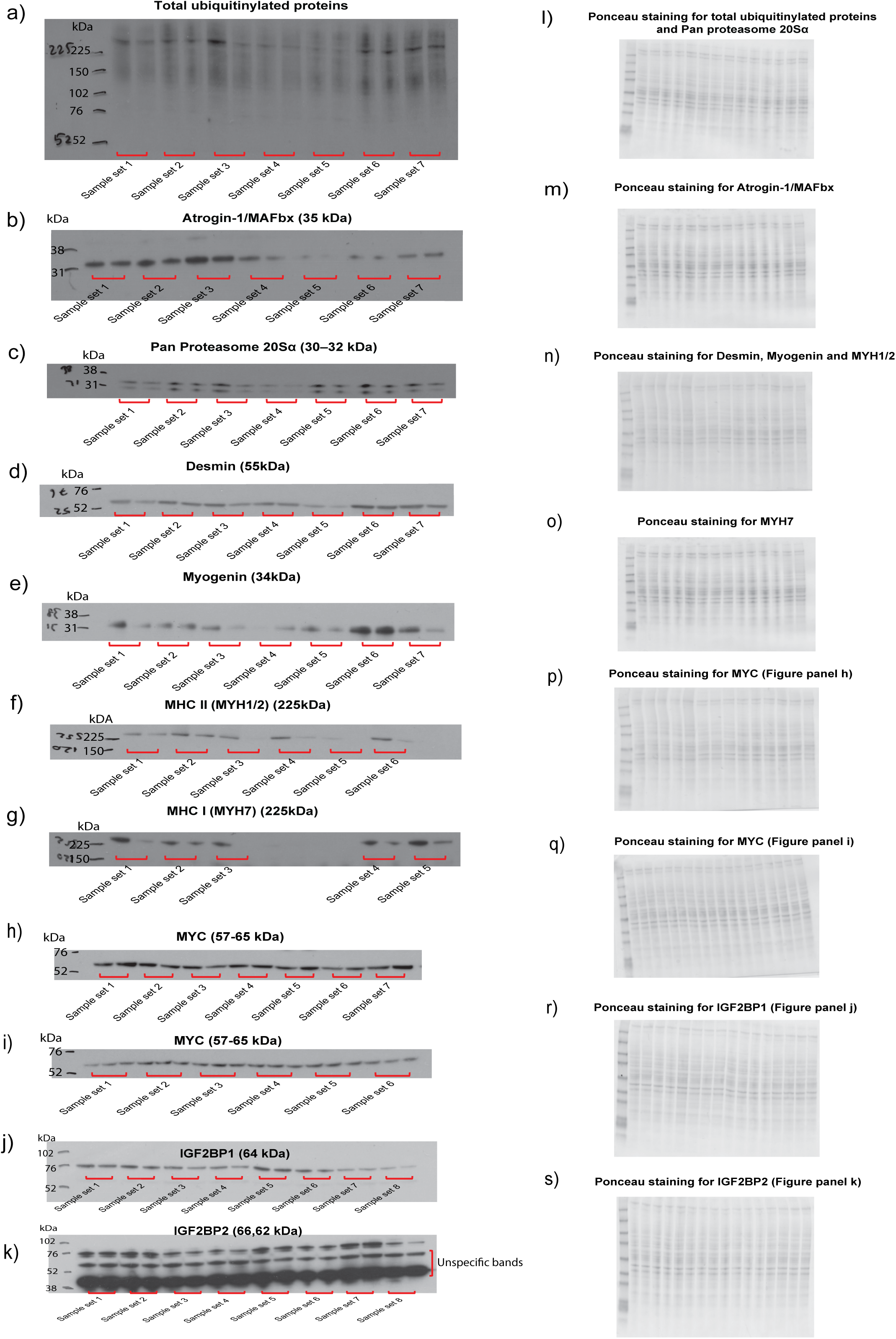
All membranes part II. Uncut membranes with the information about the sample sets/replicates for Western blot experiments of **a.** Total ubiquitinylated proteins (Fig 2c), **b.** Atrogin-1/MAFbx (Fig. 2d), **c.** PAN proteasome 20S alpha (Fig. 2e), **d.** Desmin (Extended Data Fig. 2a), **e.** Myogenin (Extended Data Fig. 2b), **f.** MYH1/2 (MyHC IIX/IIA) (Extended Data Fig. 2c), **g.** MYH7 (MyHC I) (Extended Data Fig. 2d), **h.** MYC (Extended Data Fig. 5a), **i.** MYC (Extended Data Fig. 5b), **j.** IGF2BP1 (Extended Data Fig. 6b), and **k.** IGF2BP2 (Extended Data Fig. 6c). Ponceau staining for **l.** Total ubiquitinylated proteins (Fig. 2c) and PAN proteasome 20S alpha (Fig. 2e), **m.** Atrogin-1/MAFbx (Fig. 2d), **n.** Desmin, Myogenin and MYH1/2 (MHC II) (Extended Data Fig. 2a-c), **o.** MYH7 (MHC I) (Extended Data Fig. 2d), **p.** MYC (Extended Data Fig. 5a), **q.** MYC (Extended Data Fig. 5b), **r.** IGF2BP1 (Extended Data Fig. 6b). and **t.** IGF2BP2 (Extended Data Fig. 6c). **Source data:** Source data file will be provided.

---

**Supplemetary Table 1.**
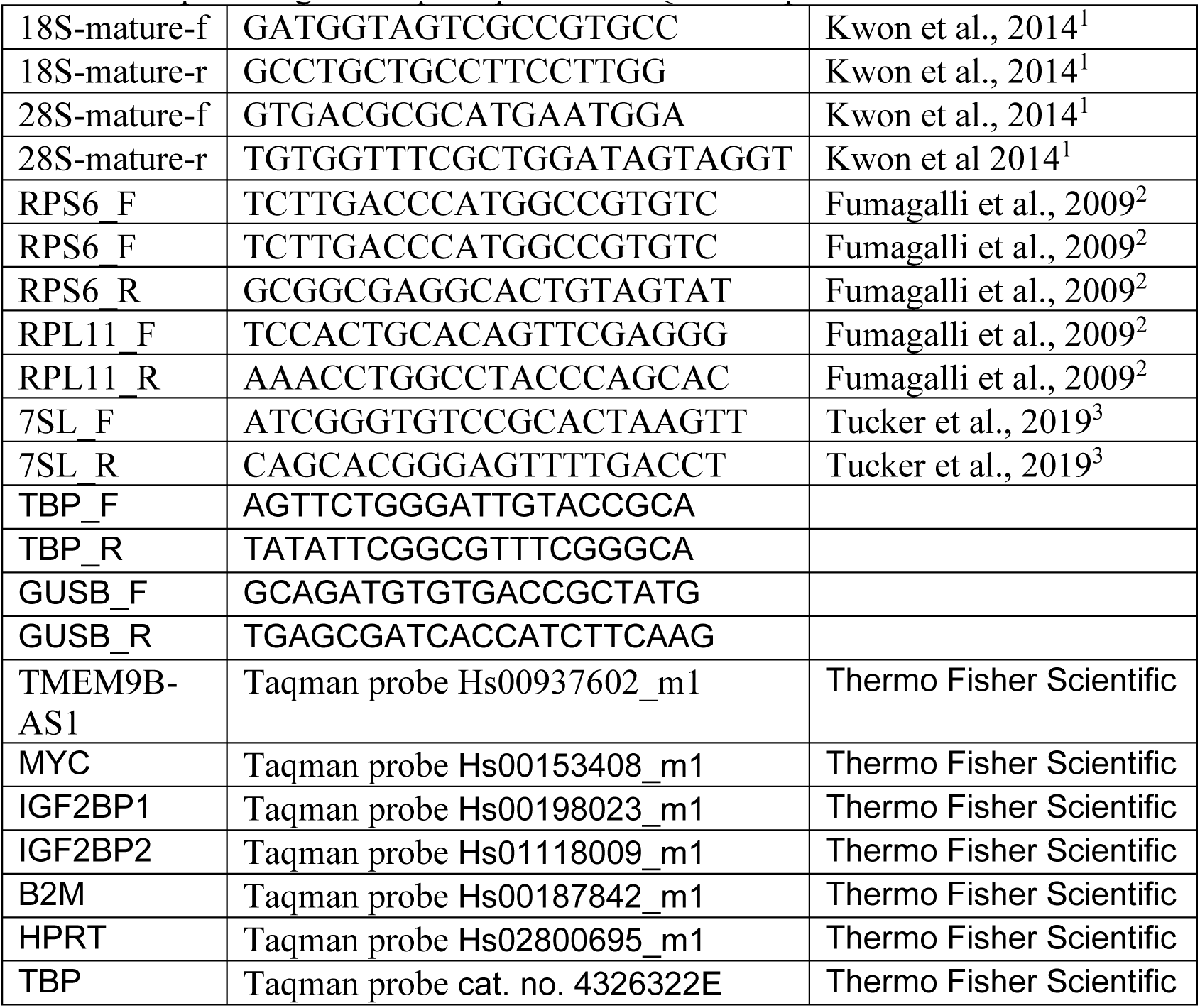
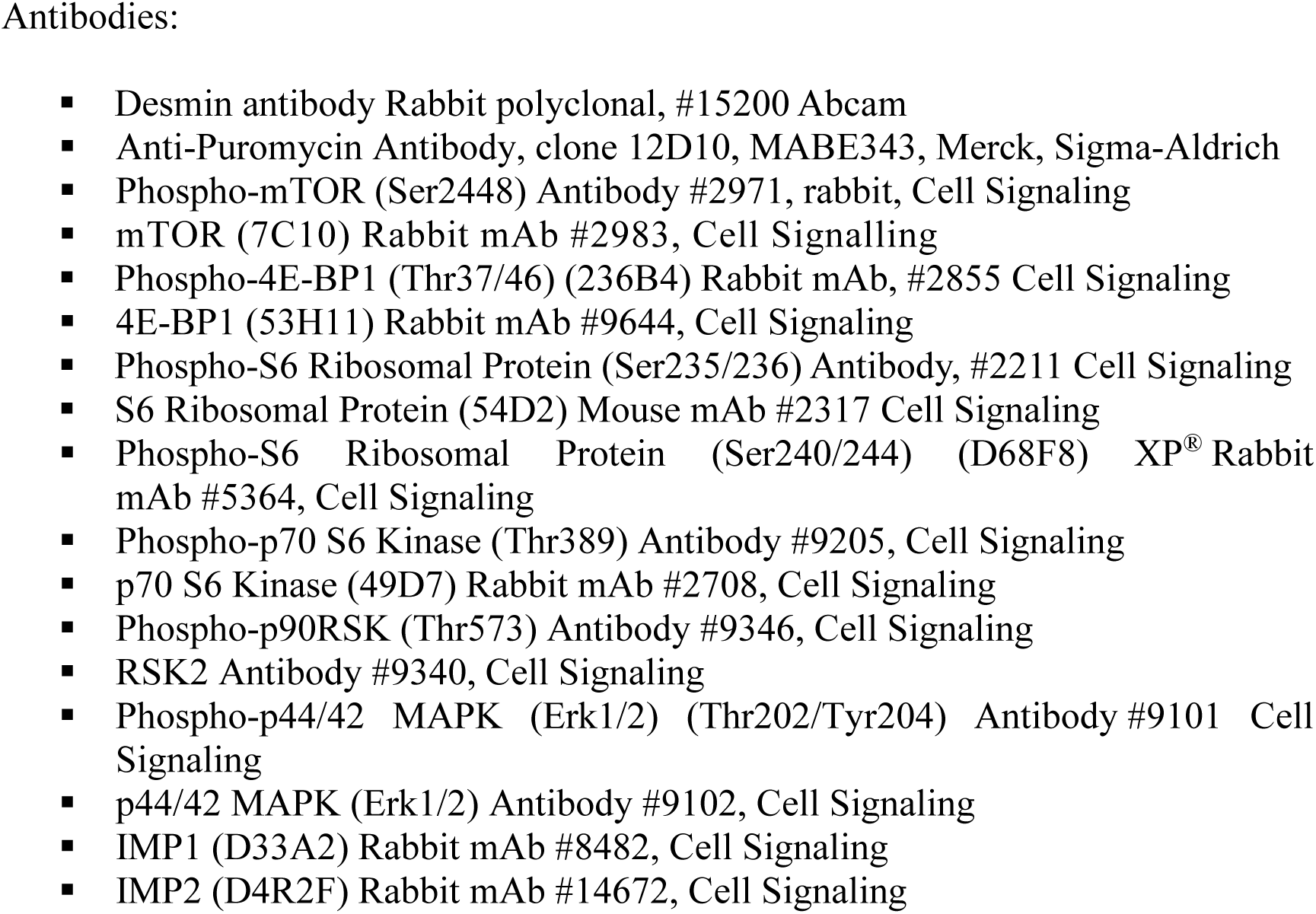
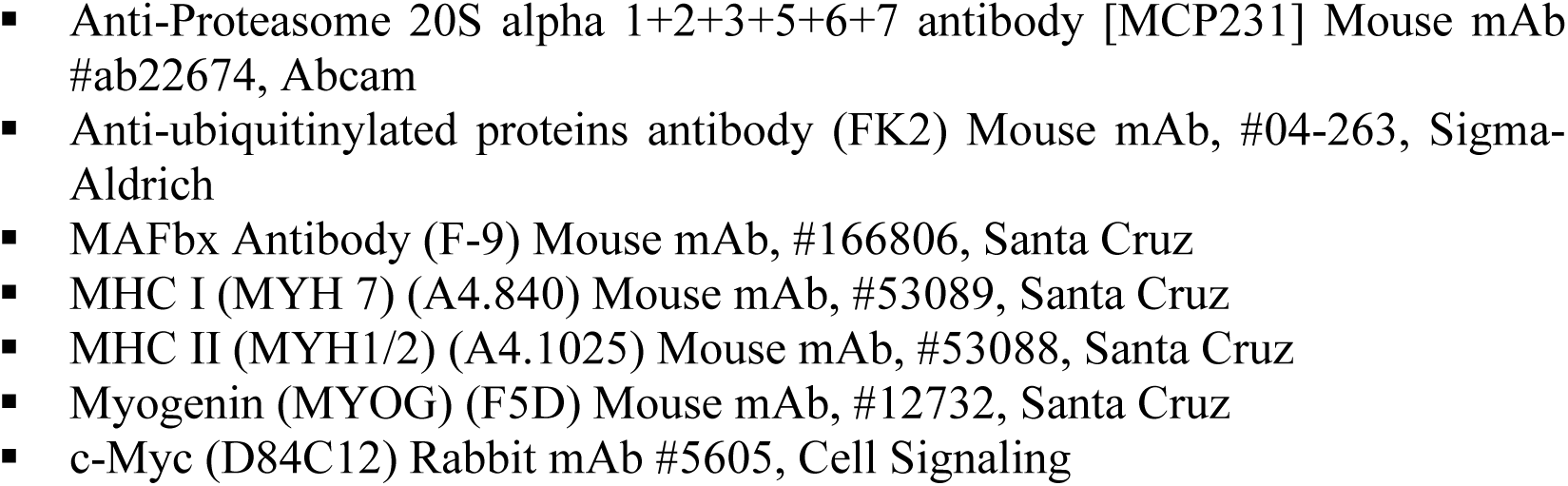
Primers and pre-designed Taqman probes for QPCR experiments:

